# Maximizing the Anti-tumor Potential of Immune Checkpoint Blockade through Modulation of Myeloid-specific CXCL16 and STAT1 Signaling

**DOI:** 10.1101/2022.03.28.485781

**Authors:** Bhavana Palakurthi, Ian H. Guldner, Xiyu Liu, Anna K. Martino, Qingfei Wang, Shaneann Fross, Ryan A. Neff, Samantha M. Golomb, Erin N. Howe, Siyuan Zhang

**Affiliations:** Department of Biological Sciences, College of Science, University of Notre Dame, Notre Dame, IN 46556, USA; Mike and Josie Harper Cancer Research Institute, University of Notre Dame, 1234 N. Notre Dame Avenue, South Bend, IN 46617, USA; Indiana University Melvin and Bren Simon Cancer Center, Indianapolis, IN 46202

## Abstract

Sensitivity to immune checkpoint blockades (ICB) depends on the overall balance of immunogenic and immunosuppressive signals in the tumor immune microenvironment (TIME). Chemotherapy as an immunostimulatory strategy showed potential in improving ICB’s clinical efficacy. Yet, evolution of highly plastic tumor-associated myeloid cells hinders ICB’s potential to reach its full therapeutic potential. In this study, we leveraged single-cell transcriptomic and trajectory analyses to delineate TIME dynamics after chemotherapy priming. We found that metronomic chemotherapy (MCT) treatment led to an accelerated T cell exhaustion through CXCL16-mediated recruitment of peripheral immature myeloid cells and expansion of STAT1-driven PD-L1 expressing myeloid cells. Inhibiting STAT1 signaling in MCT-primed breast cancer relieved T cell exhaustion and significantly enhanced the efficacy of anti-PD-1 ICB treatment. Our study leveraged single-cell analyses to dissect the dynamics of breast cancer TIME and provides a pre-clinical rationale to translate the anti-STAT1 plus anti-PD-1 combinatorial immunotherapy regimen to maximize ICB’s efficacy.

**Manuscript Summary:** Single-cell analyses on low dose chemotherapy primed breast tumor-associated immune cells demonstrates a parallel coexistence of immunogenic and immunosuppressive myeloid cell subsets. Modulating STAT1 signaling in the tumor microenvironment fine-tunes immunogenic and immunosuppressive balance and maximizes the anti-PD-1 immunotherapy efficacy in chemotherapy-primed breast cancer.

## INTRODUCTION

Cancer is a systemic disease. As such, a systemic view of cancer as an evolving ecosystem, rather than a singular view of specific cell type or gene, is essential to guide a more effective and sustained anti-cancer treatment (Hiam-Galvez et al., 2021). The divergent response to anti-cancer immunotherapy among patients highlights an urgent need for systemic interrogation of the dynamic tumor ecosystem due to the nature of immune regulation - a dynamic equilibrium, balancing the immunogenic and immunosuppressive/tolerogenic signals (Engblom et al., 2016; Galluzzi et al., 2020; Hiam-Galvez et al., 2021). Myeloid cells are prominent regulators of immune equilibrium vital to normal tissue homeostasis and tumor progression (Engblom et al., 2016). Their diverse roles in tuning tumor immune microenvironment (TIME) have been increasingly recognized as critical, sometimes decisive factors, for an optimal anti-tumor immunotherapy response (Garris et al., 2018).

Growing literature suggests a perplexing antagonistic/compensatory/redundant phenotypic plasticity among different myeloid cell subsets under various cancer contexts - from the binary M1/M2 paradigm and functionally-defined myeloid-derived suppressor cells (MDSC) to the ever growing characterization of the sub-types of myeloid cells (Engblom et al., 2016; Veglia et al., 2018). Myeloid cells are extremely heterogeneous and plastic in response to the cues imposed from TIME and systemic anti-cancer treatment (Allen et al., 2020). For instance, PARP inhibitors (in TNBC) mediate glucose and lipid reprogramming and enhance both anti- and pro-tumorigenic features of macrophages through sterol regulatory element-binding protein-1 pathway (Mehta et al., 2021). Similarly, TIME exhaustion mediated by expansion of immunosuppressive M2-like macrophages and contraction of inflammatory-macrophages during advanced renal cell carcinoma leads to decreased T cell receptor clonality (Braun et al., 2021). It is clear that the coordinated interplay between heterogeneous TIME cells dictates the anti-tumor immune responses. A systematic understanding of this coordination is critical for designing effective novel immunotherapy regimens.

Beyond the isolated view of the function of a given specific myeloid cell subset, in this study, we use breast cancer as a model to take a systematic view of tumor-associated myeloid cells in response to a commonly used chemotherapy regimen. While chemotherapy has been implemented as an immune-sensitizing strategy for ICBs, such as the TONIC trial for metastatic TNBC patients (Voorwerk et al., 2019), a significant portion of patients either do not respond or only modestly respond to ICB even after chemotherapy-priming (Galluzzi et al., 2020; Voorwerk et al., 2019). Through single-cell trajectory analysis, we observed a dynamic balance between sub-clusters of myeloids in response to chemotherapy priming. Modulating the STAT1 signaling pathway in TIME myeloid cells fine-tunes the balance between anti-cancer immunity and immunosuppression to maximize the therapeutic efficacy of anti-PD1 for breast cancer.

## RESULTS

### MCT increases T - myeloid cell colocalization, dendritic cell expansion and activity

We conducted a preclinical treatment trial to compare the impact of different chemotherapy dosing schemes (MTD or MCT) on breast tumor growth *in vivo* using three syngeneic breast tumor mouse models (Fig. S1A). In the HER2^+^ tumor model (MMTV-neu), MTD and MCT achieved similar efficacy by reducing the tumor growth to about 50% (mean relative volumes being 2 and 1.8 respectively) of the Vehicle treated tumors at the end of treatment on Day 28 (mean relative volume 3.7) (Fig. 1A, left). In the triple negative breast cancer (TNBC) mouse model (C3-1-TAg), MTD did not, but MCT did significantly reduce the tumor growth to 46% of Vehicle treated tumors (Fig. 1A, center). In the adenocarcinoma tumor model (MMTV-PyMT), MCT significantly reduced the tumor growth to 50% of Vehicle treated tumors (Fig. 1A, right). In MMTV-neu mice, MTD extended the (tumor volume doubling time) progression-free survival (PFS) from ∼13 to 21 days, while MCT prolonged the PFS from 13 to 27 days (Fig. 1B, left). In C3-1-TAg mice, both MCT and MTD extended the PFS from 6 to ∼ 14 days (Fig. 1B, center). In MMTV-PyMT, MCT doubled the PFS from 14 to 28 days (Fig. 1B, right). Moreover, tumor proliferation, as depicted by H-Score of Ki-67 staining, decreased 1.8-2.6 folds compared to respective Vehicle control after either MTD or MCT treatment (Fig. S1B).

**Figure 1.**
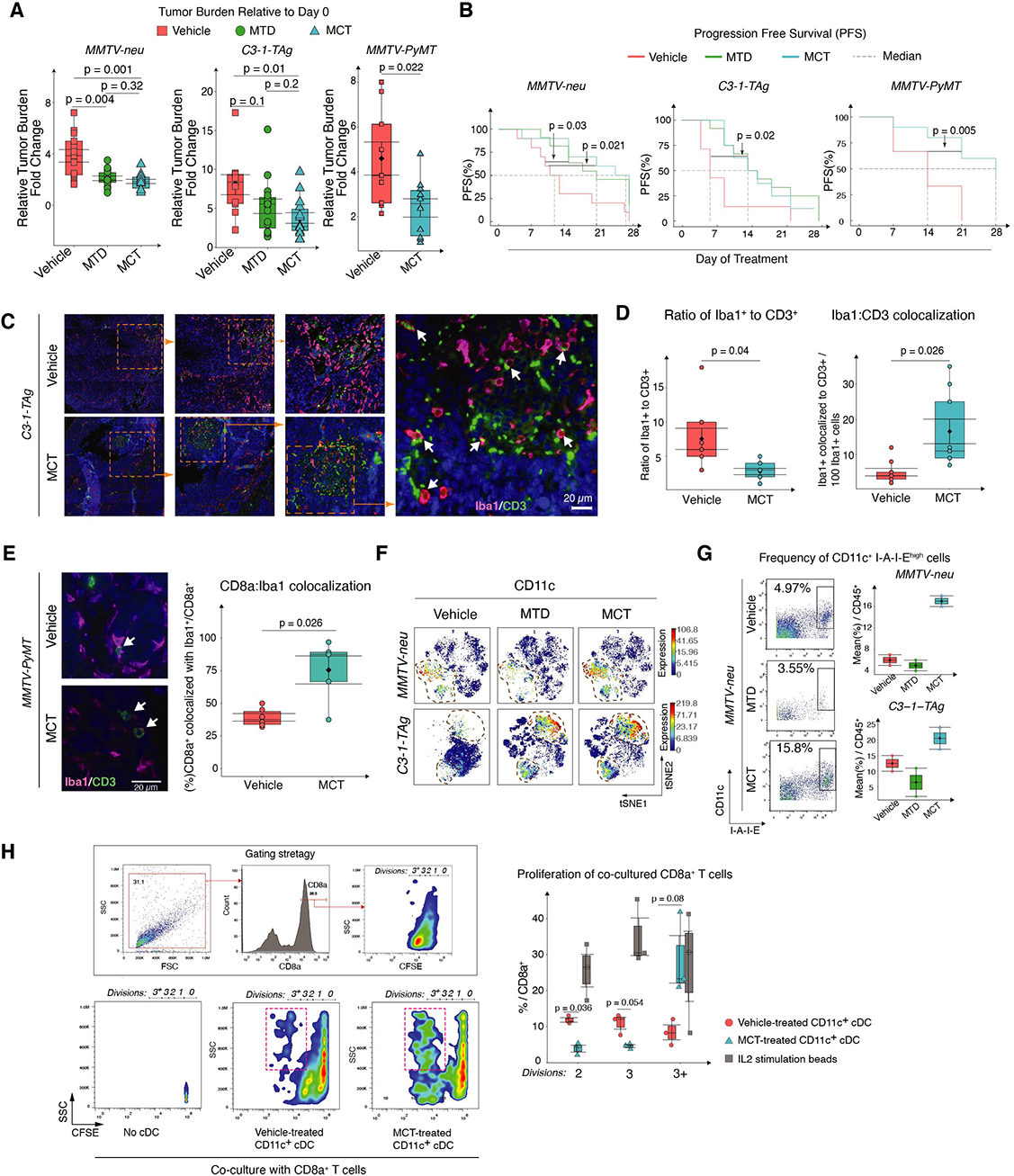
MCT increases T - myeloid cell colocalization, dendritic cell expansion and activity in breast tumor. (A) Box plots show tumor burden on Day 28 relative to Day 0 of treatment in MMTV-neu (left), C3-1-TAg (center), and MMTV-PyMT (right) mice. (B) PFS (%) indicates the tumor volume doubling time in MMTV-neu (left), C3-1-TAg (center), and MMTV-PyMT (right) (generated using the Survimer R package, log rank p-value). (A-B) 3 to 5 mice pooled over three cohorts (n = 10- 13 per group). (C) Representative IF images of C3-1-TAg breast tumor tissues stained with Iba1, CD3, and DAPI. Inset (far right) shows the CD3+ and Iba1+ cell colocalization indicated by arrows. (D) Box plots quantify Iba1+ and CD3+ cell ratio (left) and their colocalization (right) (n = 9 slides per group). (E) Representative IF images of MMTV-PyMT breast tumor tissues stained with Iba1, CD8, and DAPI. Box plots quantify Iba1+ and CD8+ cell colocalization (n ≥ 3). (F) CyTOF data t-SNE plots compare CD11c expression in MMTV-neu and C3-1-TAg tumors upon treatment with either Vehicle, MTD, or MCT. (G) Biaxial plots (left) compare and box plots (right) quantify the mean % of CD11c+ I-A-I-E+ cells among all the CD45+ cells identified using CyTOF. (H) Gating strategy (left) compares CFSE levels on CD8a+ T cells with different division frequencies in co-culture with CD45+ CD11c+ cells isolated from C3-1-TAg tumors. Box plot quantifies in vitro proliferating T cells (%) with different division frequencies when co-cultured with CD11c+ cells (n = 3). Data in box plots represent Mean ± SEM, Diamonds Means, Black lines Medians, and T-test p-values.

Above results derived from three breast cancer models are consistent with observed clinical benefit of MCT (Pasquier et al., 2010; Pogoda et al., 2017). Given the perceived myeloid toxicity and potential benefit of MCT in modulating TIME, we sought to understand the immune cell composition and spatial co-localizations within the tumor under different treatment conditions. After treating non-tumor bearing C3-1-TAg mice either with Vehicle, MTD, or MCT respectively for one week, the count of CD45^+^ cells in peripheral blood decreased from 5*10^3^ in Vehicle treated to 3*10^3^ per µL in MTD mice while their count increased to 14*10^3^ per µL of blood in MCT-treated mice (Fig. S1C). However, in the tumor bearing mice, all treatments significantly reduced the CD45^+^ immune cell percent in the peripheral blood (Fig. S1D), suggesting tumor formation profoundly changed systemic immune regulation (Allen et al., 2020; Lucas et al., 2013). Interestingly, MCT-treated tumors showed a significant increase of tumor-infiltrating CD3^+^ lymphocytes (TILs) (Fig. S1E) with evident “hotspot” areas co-localized with Iba1^+^myeloid cell and CD3^+^TILs (Fig. 1C-D), suggesting an increased spatial proximity between those cells. The ratio of tumor-infiltrating myeloid cell to T cell ratio decreased with MCT treatment (Fig. 1D, left). Importantly, the percent of myeloid cells colocalizing with T cells significantly increased in MCT-treated tumors (Fig. 1D, right). Similarly, in the MMTV-PyMT model, MCT treatment preserved the CD45^+^ TILs (Fig. S1F) and CD8a^+^ T cells that co-localized with Iba1^+^ myeloid cells increased in MCT-treated MMTV-PyMT tumors (Fig. 1E).

To examine the MCT-mediated tumor-associated immune landscape modulation, we immunophenotyped primary breast tumors under Vehicle, MTD, and MCT strategies using a 19-antibody CyTOF panel (**Supplemental Table 1**). CyTOF results confirmed that MCT preserves TIL (Fig. S1G). Next, we downsampled CyTOF detected immune cells in each treatment group to the equal total number per treatment group and clustered them using the viSNE algorithm (Fig. S1H). After pooling the data from two independent CyTOF experiments, we quantified the changes of tumor-associated immune cell subtypes based on traditional manual gating (Fig. S1I). Interestingly, CD11c^+^ subset constituted one third of the whole myeloid cell compartment within MCT tumors (Fig. 1F). When quantified based on manual gating, the proportion of CD11c^+^ I-A-I-E^+^, representing the functionally activated conventional dendritic cells (cDCs), among other immune cells increased in MCT treated tumors compared to Vehicle and MTD treated groups (Fig. 1G).

To examine the functional impact of TIL cDCs on CD8^+^ T activity after MCT treatment, we performed an in *vitro* T cell proliferation assay by co-culturing CD11c^+^ TIL cells with naive T cells. CD45^+^CD11c^+^ cDCs sorted from TNBC tumors stimulated the expansion of naive CD8^+^ T cells as shown by the CFSE dye dilution (Fig. 1H). Importantly, co-culture with MCT-treated CD11c+ cDC led to a significant increase of proliferative T cells which have divided more than three times (8.4% and 28.7% in Vehicle and MCT respectively) with a higher granularity (red boxes shown in biaxials) (Fig. 1H), indicating higher activation status.

### CITE-seq reveals distinct myeloid subsets co-existed in MCT-treated TME

To further characterize the immune cell transcriptome changes after MCT treatment at single-cell level, we employed Cellular Indexing of Transcriptomes and Epitopes by Sequencing (CITE-seq) with a 29-antibody panel representing well-known immune cell surface markers (**Supplementary Table 2**, immune Cell-ID). Cell-ID based gating using CITE_CD45 antibody and high RNA expression of *Ptprc* gene identified a total of 542 high quality CD45^+^ TILs per treatment group from quality controlled immune cell singlets. We manually gated these CD45^+^ cells into myeloid (CD11b^+^) and lymphoid (CD3^+^) cell subsets (Fig. S2A). Among the CD11b^+^ myeloid cells, we identified cDCs and macrophages based on known markers that have been previously identified in human TILs (Binnewies et al., 2019; Broz et al., 2014; Laoui et al., 2016), including tumor-associated dendritic cells – CD11b^+^ cDC2s, CD103^+^ cDC1s, and Monocytic DCs (together abbreviated as TADs); and CD11b^+^ or CD11c^+^ tumor-associated macrophages (together abbreviated as TAMs) (Fig. S2A). RNA-based Uniform Manifold Approximation and Projection (UMAP) analysis revealed 17 different single cell RNA clusters - here we refer to them as Seurat RNA Clusters (Fig. 2A, left). Broadly, these clusters exhibited known transcriptional signatures representing unique immune cell types (as labeled below UMAP and **Supplementary Table 3**). Interestingly, canonical TADs and TAMs identified conventional surface makers (immune Cell-ID) gating (Fig. S2A) did not overlap with any specific RNA clusters when projected on the RNA UMAP (Fig. 2A, right). This discordance between the cell surface and transcriptional status suggested a high level of transcriptional plasticity of myeloid cells (Maier et al., 2020; Satpathy et al., 2012).

**Figure 2.**
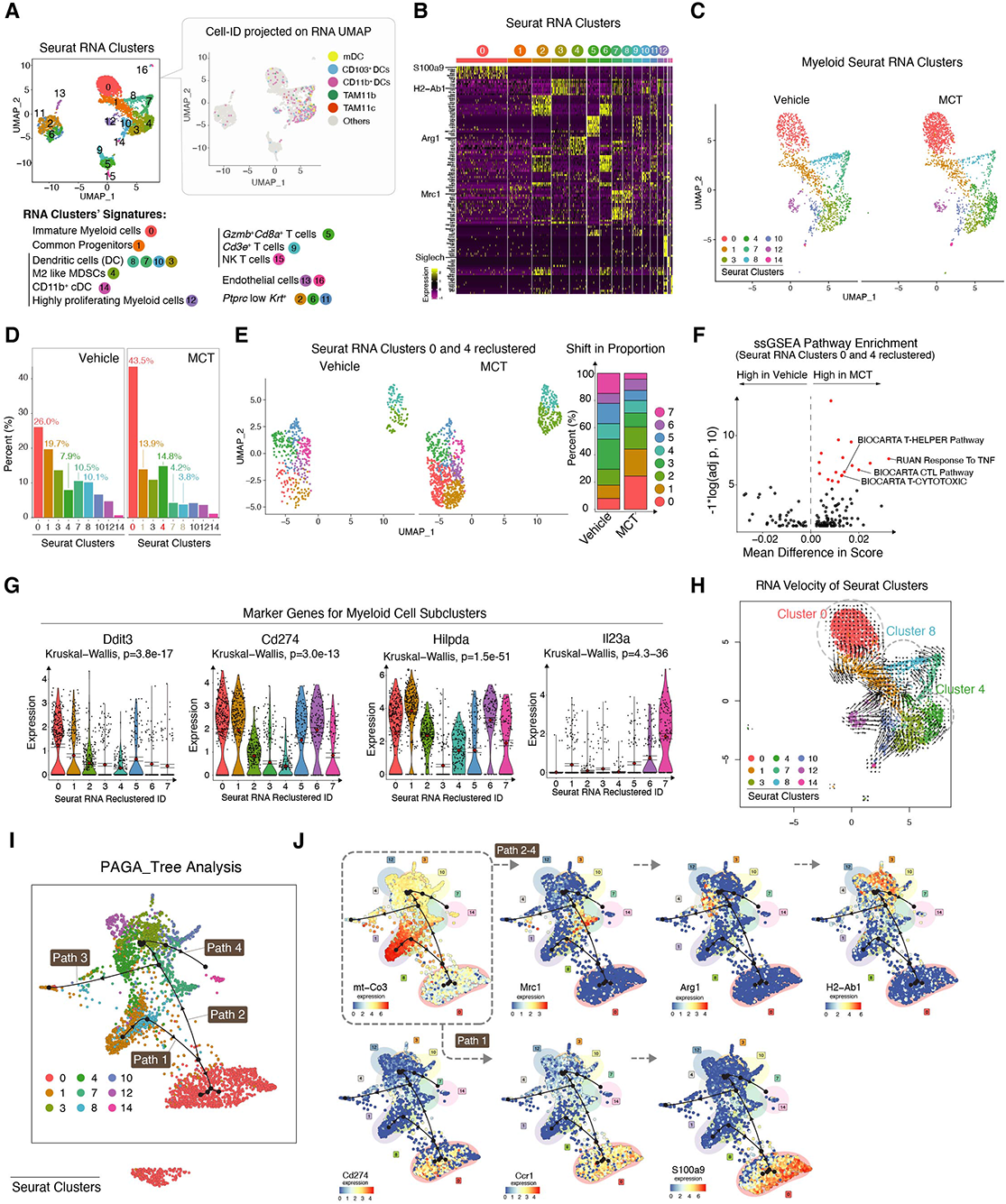
CITE-seq reveals distinct myeloid subsets co-exist in MCT-treated TME. (A) UMAP clusters of TNBC derived single cells: RNA defined clusters (left) and Cell-ID defined clusters of TADCs and TAMs (right) (in both the UMAPs: dots = cells, Cell-ID UMAP: colored dots = TADs and TAMs, and gray dots = non TADCs and non TAMs). (B) Heatmap shows expression of top differentially expressed genes (DEGs) among RNA Seurat Clusters (C) UMAP clusters of transcriptionally defined myeloid cells split by treatment. (D) Bar plot shows (%) proportion of myeloid Seurat Clusters projected in (C). (E) UMAP (left) split by treatment projects reclustered Seurat Clusters 0 and 4. Stacked bar plot (right) shows shifts in the proportion of different clusters identified after reclustering. (F) Volcano plot of differentially expressed gene pathways (DEGPs) between MCT and Vehicle treated clusters in (E). (G) Violin plots show expression of marker genes of clusters from (E). (H) UMAP projects RNA velocity (length of arrow tail) of myeloid Seurat Clusters. (I) PAGA tree predicts trajectory paths of myeloid Seurat Clusters. (J) PAGA trajectory tree projects marker genes of trajectory clusters 7, 4, and 3 in Paths 2-4 and Cluster 0,1 in Path 1. In A to J, n = cells pooled from 3 biological repeats per treatment group.

Next, we examined marker genes of each RNA cluster identified (Fig. 2B and **Supplementary Table 3**). Cluster 0 expressed inflammatory (*S100a8/9*), immunosuppressive genes (*Hcar2* and *PD-L1*), *Ccr1*, *Nfkbia*, and *Nlrp3*. Since these cells lacked DC maturation genes and were expressing *Nlrp3*, an immature DC marker (Ghiringhelli et al., 2009), we categorized Cluster 0 as immature myeloid cells. Cluster 1 expressed DC progenitor genes (*Trps1*, *Rel*, and *Etv3*). Clusters 2, 6, and 11 expressed Keratin genes (*Krt7, Krt8, and Krt18*), suggesting their potential tumor origin. Clusters 3, 7, 8, and 10 gene signatures overlapped with both DC and macrophage gene signatures (*Itgax*, *Csf1r*, *Irf8*, *Mrc1*, *Etv3*, *Cd74*, *H2-Ab1*, and *Cd86*). Cluster 4 expressed MDSC gene signature (*Arg1, Irf8*, *Itgam,* and *Lgals3*). Cluster 12 expressed high proliferation and myeloid specific genes (*ki-67*, *Cd86*, and *Cd72*). Clusters 13 and 16 expressed fibroblastic and endothelial genes (*Col1a2, Col1a1, Fgf7, and Fgf2*). Cluster 14 comprises cells with classic cDC gene signature (*Btla4*, *Siglech*, *Flt3l*, P2ry14, and *Irf8*). Clusters 5, 9, and 15 expressed lymphoid specific gene signature (*Cd3e*, *Cd3g*, and *Cd3d*). Cluster 5 hosted *Gzmb*^+^ *Cd8a*^+^ T cells and 15 hosted *Nk* T cells that expressed *Fasl* and *Gzmb*. When examining single cell transcriptome marker genes of each Cell-ID based TADs and TAMs cluster identified, all the TADs and TAMs showed significant similarities in their transcriptional profile (Fig. S2B). We then examined the overall gene expression modulation within TADs and TAMs under MCT. After MCT treatment, TADs and TAMs expressed a higher level of genes related to T cell dysregulation (*Lgals3*), immune cell accumulation (*Fth1*), and energy production (*Pgam1*) (Fig. S2C).

Among the different immune cell types, myeloid clusters (transcriptionally identified) constituted nine unique Seurat Clusters -Clusters 0, 1, 3, 4, 7, 8, 10, 12, and 14 (Fig. 2C). Clusters 0 and 4 expanded from 26% and 7.9% in Vehicle to 43.5% and 14.8% in MCT respectively upon MCT treatment, whereas Clusters 1, 7, and 8 depleted from 19.7%, 10.5%, and 10.1% in Vehicle to 13.9%, 4.2%, and 3.8% in MCT (Fig. 2D). The MCT treatment led to downregulation of mitochondrial genes (*mt-Cytb*, *mt-Nd*2, and *mt-Nd*4) and up-regulation of *Fth1*, *Cstb*, *Hilpda*, and *Cd*274 in myeloid cells (Fig. S2D). Gene set enrichment analysis showed up-regulation of ribosome, protein chain elongation, and *TGFβ* pathways in MCT-treated myeloid cells (Fig. S2E and **Supplementary Data Table 4**). Next, we subsetted two clusters (Clusters 0 and 4) that were significantly changed after MCT and reclustered them to learn their identity and heterogeneity. We found eight unique transcriptional UMAP clusters (Fig. 2E, left and **Supplementary Table 5**) within subsetted Seurat RNA Clusters 0 and 4. Among these eight clusters, Clusters 0, 1 expanded while Clusters 7 proportions contracted after MCT (Fig. 2E, right). ssGSEA Pathway Enrichment analysis further showed an enriched T cell helper, cytotoxic, and tumor necrosis factor (TNF) pathways after MCT treatment (Fig. 2F and **Supplementary Table 6**). Interestingly, Clusters 0 and 1 cells were marked by high expression of genes related to the immune system’s response to inflammation (*Ddit3)* (Mogilenko et al., 2019), immune checkpoint (*Cd274*), hypoxia-induced anti-inflammatory mitochondrial metabolism process in myeloid cells (*Hilpda*) (DeBerge et al., 2021). Cluster 7 cells express a high level of Il23a, resembling the gene signature of myeloid suppressor cells (MSDC) observed in castration-resistant prostate cancer (Calcinotto et al., 2018)(Fig. 2G **and** Fig. S2F).

Coupling the RNA velocity (La Manno et al., 2018) (Fig. 2H) and the PAGA tree trajectory (Saelens et al., 2019) analyses (Fig. 2I,J **and** Fig. S2G), we further examined the evolution trajectories (cellular status transition) of myeloid Seurat RNA Clusters. Interestingly, despite the fact that Cluster 0 and 4 are the myeloid clusters which expanded the most in MCT tumors, Clusters 0, 4, and 8 had the lowest RNA velocity as depicted by the length of the arrow’s tails. This suggests that these clusters are the three major transcriptome anchor points of myeloid cells co-existing in the tumor microenvironment (Fig. 2H, encircled clusters). PAGA tree analysis further delineated the specific trajectory paths that lead to the convergence of various myeloid cells into the Clusters 0, 4, and 8 (Fig. 2I). Cluster 1 (Common Progenitors) had two main future states or trajectory paths - Path 1 and Path 2 (Fig. 2I, arrow heads). Through Cluster 8, Cluster 1 evolved along Path 1 into Cluster 0 (immature MDSC-like cells expressing Cd274, Ccr1, and S100a9) or bifurcated into Path 2 (Fig. 2J and Fig. S2G). Along the trajectory Path 2, Cluster 7 cells (*Mrc1*+) emerged and then evolved into Cluster 3 (*H2-Ab1* and *Cd86* high DCs) and Cluster 4 (*Arg1*+ M2-like MDSCs) (Fig. 2J and Fig. S2G). The Path 3 was an offshoot of Path 2 that then returned into a sub-portion of Cluster 1. Cluster 10 (cDC) continued evolving into *H2-Ab1*-expressing Cluster 14 (cDC2) via Path 4 (Fig. 2J). Collectively, our data revealed that tumor infiltrating myeloid cells after MCT treatment showed highly heterogeneous transcriptome status, balancing the immune stimulatory and immunosuppressive environment at TME.

### Mapping tumor-associated myeloid cell transcriptome regulatory networks

MCT treatment led to an increase of Cluster 0 cells stopping at the end of trajectory Path 1, suggesting an intermediate cellular status between myeloid progenitors (Cluster 1 and 8) and fully differentiated lineages (Clusters 3, 7, 10, 12, and 14). First, to characterize the identity of Cluster 0 cells, we compared their gene signature with the marker genes of MDSCs (Veglia et al., 2018), M1/M2 macrophages (Xue et al., 2014), Monocyte-derived DCs (Tang-Huau et al., 2018), Mature DCs, and regulatory DCs (Maier et al., 2020) (Fig. 3A). Cluster 0 cells showed minimal transcriptional overlap with above known tumor associated myeloid cells types, except overexpressing inflammatory response genes *S100a8/9* and myeloid differentiating gene *Cebpb*, and pathogen recognizing gene *Cd14* (Fig. 3A **and** Fig. S3A-H). However, classic MDSC marker genes such as *Ly6c*, *Ly6g*, and *Arg1* were not expressed by Cluster 0 cells. Additionally, *Ccr1* inflammatory response gene (expressed by a subset of MDSCs (Inamoto et al., 2016; Li et al., 2019) was differentially expressed by Cluster 0 cells. Next, to explore the transcriptional programs that drive specific myeloid cell cluster’s evolutionary trajectory, we performed single-cell regulatory network inference and clustering (SCENIC, detailed in Methods, Fig. 3B Schematic) (Aibar et al., 2017). In brief, a Transcription Factor (TF) and its predicted target motifs co-expressed in a single-cell are identified as a Regulon. The enrichment scores of the Direct target motifs are normalized and scored as Normalized Enrichment Scores (NES). The activity of each regulon is scored as either active or inactive which is then used to calculate the activity percentage (% of cells with an active regulon) and cluster cells to identify cluster specific regulatory networks (Fig. 3B). We identified multiple active regulons either generally enriched to all the myeloid clusters or specific to one cluster or a subset of clusters (Fig. 3C). Notably, *Nfkb1*, known to regulate immunological responses to infections and diseases, was active in all the clusters. Cluster 0 specifically had highly active *Cebpb*, *Bhlhe40*, and *Atf3* that regulate cellular growth, differentiation, and immune response during stress. On the other hand, *Bcl11a* essential for plasmacytoid dendritic cell differentiation (Ippolito et al., 2014) was highly specific to Cluster 14. Interferon signal regulating gene *Irf8* along with general cellular growth regulators such as *Fos*, *Egr1*, and *Jun* were active in all the clusters with intermediate cellular state - Clusters 12, 3, 4, 10, 1, 7, and 8. Interestingly, we observed an opposite trend with *Foxp1* and *Fosl2* activity aross the clusters. While *Foxp1* which is upregulated upon DC maturation was highly active in certain clusters including Cluster 14, where *Fosl2* activity was lower and vice versa (Fig. 3C, lower portion). We then projected some of these regulons with activity specific to a trajectory path/cluster on UMAP (Fig. 3D). Clearly, Cluster 0 hosted cells with highest *Cebpb* and *Fosl2* (Sarode et al., 2020) activity specific to PAGA tree trajectory Path 1 while Cluster 14 hosted cells with *Bcl11a* activity. Similarly, *Foxp1* and *Irf8* were specific to trajectory Paths 2 and 4.

**Figure 3.**
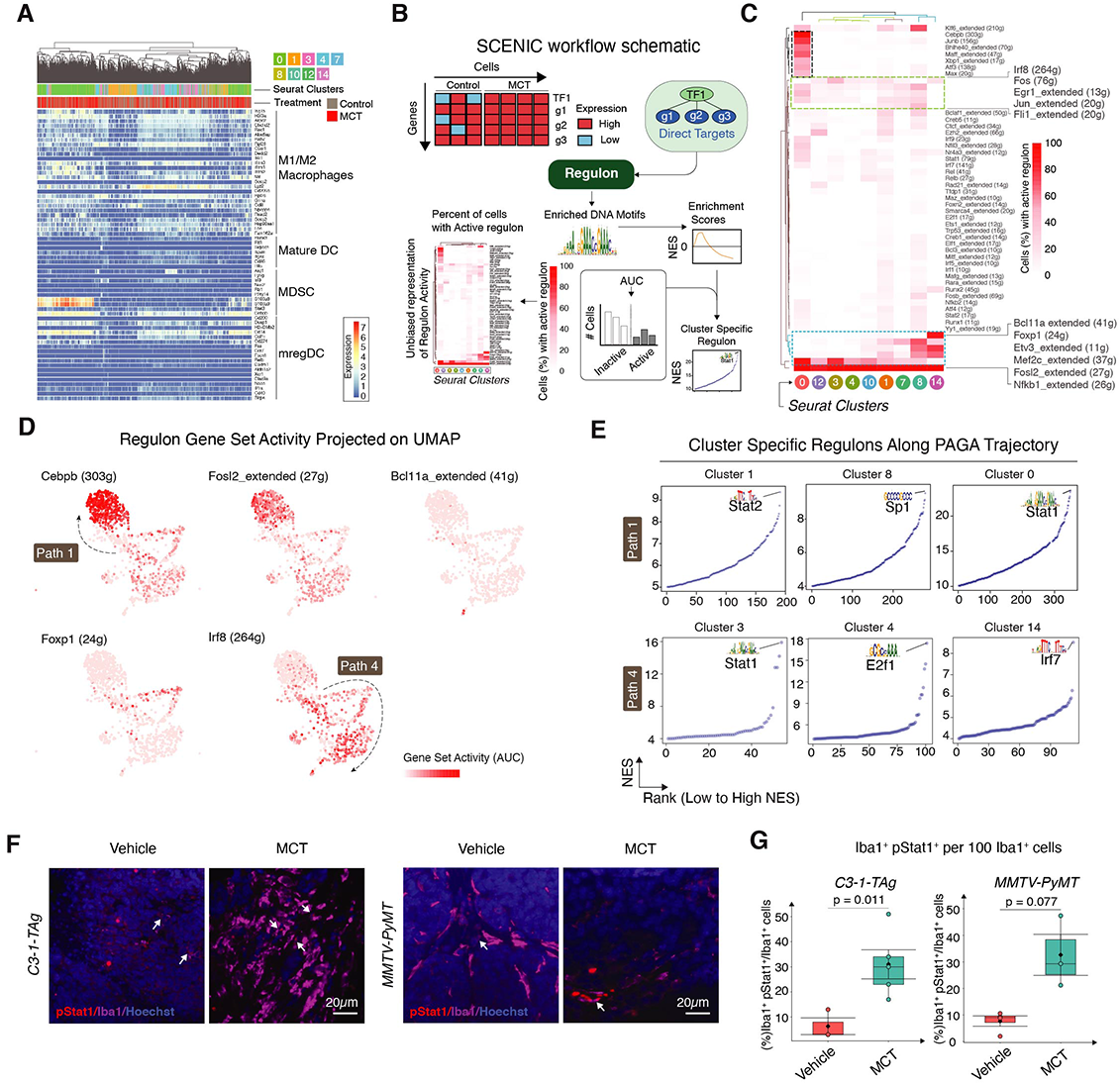
Mapping tumor-associated myeloid cell transcriptome regulatory networks using Single-Cell regulatory Network Inference and Clustering (SCENIC) (A) Heatmap shows expression of indicated cell type marker genes by myeloid Seurat clusters. (B) Schematic shows SCENIC workflow. (C) Heatmap shows percent of cells in a cluster with active regulon. (D) UMAP projects the regulon gene set activity. (E) Scatter plots show regulons specific to (top row) Clusters 1, 8, and 0 (left to right), (bottom row) Clusters 3, 4, and 14 (left to right) and their NES. Each dot is a representative of one NES (see Methods). (F) Representative IF images of murine C3-1-TAg tissues stained with pStat1 (Ser 727), Iba1, and CD3 and MMTV-PyMT tissues with pStat1(Ser 727) and Iba1. (G) Box plot shows quantification of pStat1+ Iba1+ cells infiltrating the C3-1-TAg and MMTV-PyMT tumor (Vehicle and MCT-treated, n ≥ 3). Data in box plots represent Mean ± SEM, Diamonds Means, Black lines Medians, and T-test p-values.

To dissect the regulon specificity for each of the clusters related to trajectory Path 1 and 4, we identified the specific regulons for each of the Seurat clusters along the trajectory Path 1 and 4 (Fig. 3E and **Supplementary Table 7**). *Stat2*, *Sp1*, and *Stat1* scored the highest NES of 9.36, 9.67, and 23.6 in Clusters 1, 8, and 0 respectively. *Stat1*, *E2f1*, and *Irf7* scored the highest NES of 15.9, 17.4, and 8.52 in Clusters 3, 4, and 14 respectively (Fig. 3E). To validate the predicted activation status of *STAT1* regulon after MCT treatment, we co-stained C3-1-TAg and MMTV-PyMT tumors with Iba1 (myeloid marker) and pStat1 (Ser727) - an activation marker for *STAT1* (Decker and Kovarik, 2000) (Fig. 3F). MCT treatment expanded the Iba1^+^pStat1^+^ cells (white arrows) from a mean of 6 and 8% (Vehicle treated) to 31 and 33% of all the Iba1^+^ cells in C3-1-TAg (left) and MMTV-PyMT (right) respectively (Fig. 3G).

### MCT-induced CXCL16 mediates intratumoral immune dynamics and ICB efficacy

As Path 4 comprises diverse myeloid cell clusters, we questioned whether these myeloid clusters share any common changes upon MCT treatment. First, we grouped cells from Path 4 clusters (Seurat Cluster 3, 7, 8, and 10) and compared marker genes to Cluster 0 cells (Fig. 4A). Mitochondrial genes (*mt-Co1*), DC maturation genes (*Cd83*), and hemoglobin genes (*Hba-a1*) were highly upregulated in Path 4 clusters while immune response modulator (*Fth1*), immune checkpoint protein (*Cd274*), and anti-inflammatory gene (*Hilpda*) were higher in Cluster 0. In addition, several inflammation regulatory cytokines were differentially expressed. *Cxcl12* and *Ccl3* were upregulated in Cluster 0 cells and *Ccl2*, *Cxcl10*, and *Cxcl16* were upregulated in Path 4 clusters (Fig. 4A). Among these cytokine related genes, the expression of *Cxcl2*, *Ccl2*, *Cxcl10*, and *Ccl3* did not differ significantly upon MCT treatment (Fig. S4A). However, the Path 4 cluster-specific *Cxcl16* expression levels were significantly up-regulated in MCT-treated Path 4 clusters (Fig. 4B, left and Fig. S4B). *Cxcl16* levels are known to increase in exhausted tissue microenvironments and are correlated with high expression of interferon gamma (Veinotte et al., 2016). In line with this notion, IFN receptor (*Ifnγr1* and *Ifnγr2*) expression levels were higher in Path 4 clusters cells upon MCT treatment (Fig. 4B, right).

**Figure 4.**
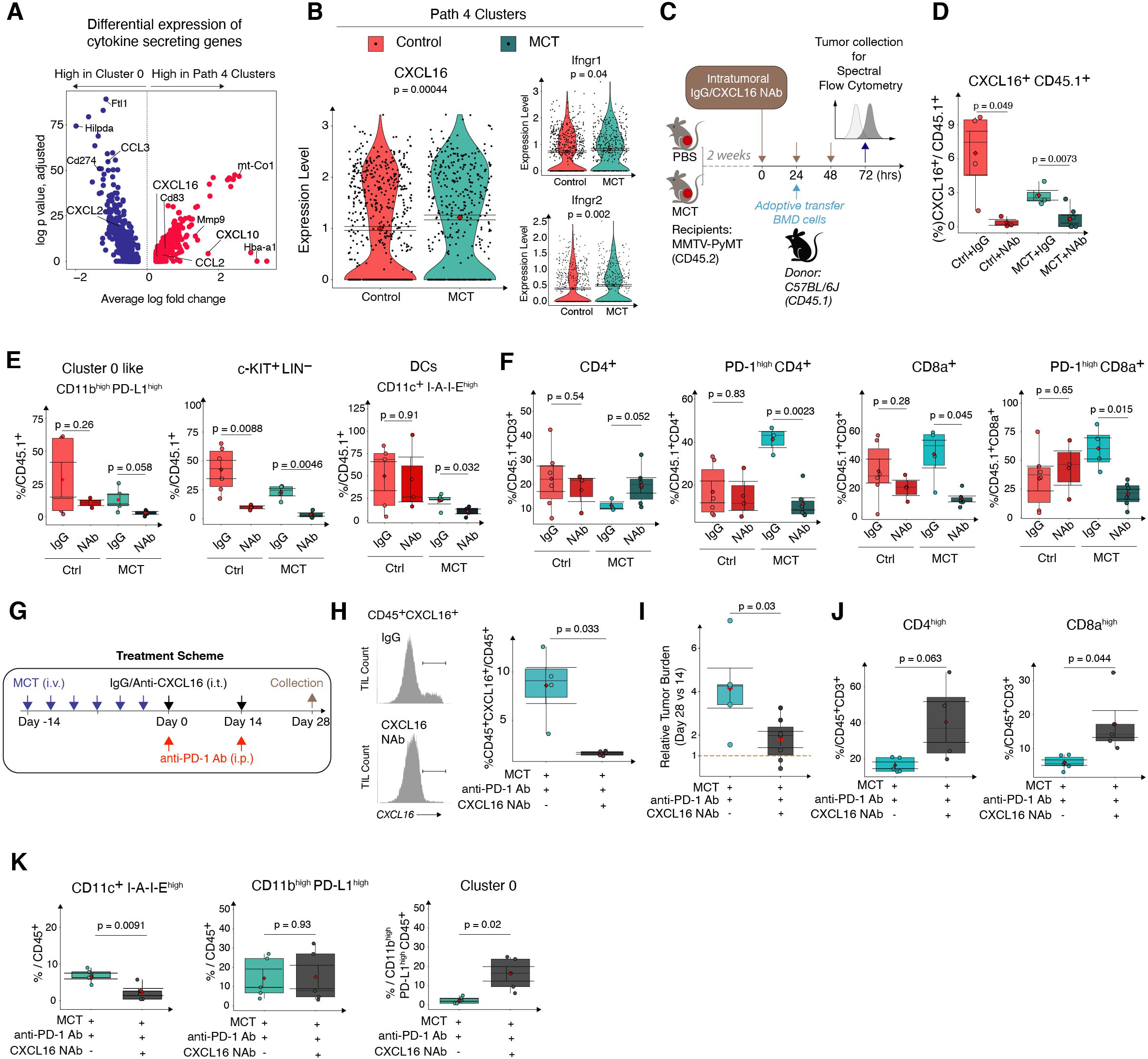
MCT-induced CXCL16 mediates intratumoral immune dynamics and ICB efficacy. (A) Volcano plot shows genes expressed differentially between Cluster 0 and DC clusters. (B) Violin plots show the expression of CXCL16, IFN-γr1, and IFN-γr2 by the DC clusters. (C) Schematic shows the intratumoral IgG or CXCL16 NAb injection into CD45.1^+^ cell adoptive transfer recipients treated with either PBS or MCT. (D-E) Box plots show the infiltration of CXCL16^+^ CD45.1^+^ cells, CD11b^high^PD-L1^high^, c-KIT^+^LIN^-^, and CD11c^high^I-A-I-E^high^ cells in PBS and MCT-treated tumors upon CXCL16 NAb injection. (F) Box plots show changes in CD4^+^, PD-1^high^ CD4^+^, CD8a^+^, and PD-1^high^ CD8a^+^ T cells in PBS and MCT-treated tumors upon CXCL16 NAb injection. (G) Schematic shows combinatorial treatment with anti-PD-1 antibody and CXCL16 NAb in C3-1-TAg mice. (H) Histograms (left) show CXCL16^+^ CD45^+^ cell frequency and box plot (right) quantify CXCL16^+^ CD45^+^ cells in different treatment conditions. Box plots quantify (I) the tumor burden on Day 28 relative to Day 14, (J) CD4 and CD8a T cells associated with tumors, and (K) CD11c^high^I-A-I-E^high^, CD11b^high^PD-L1^high^, and Cluster 0 cells in different treatment conditions.

To elucidate the functional impact of CXCL16^+^ cells on activating T cells, we performed an in *vitro* proliferation assay by co-culturing CXCL16^+^ cells with naive T cells. CXCL16^+^ cells enriched from TNBC tumors stimulated the expansion of naive both CD8^+^ T and CD4^+^ T cells as shown by the CFSE dye dilution (Fig. S4C). Importantly, co-culture with MCT-treated CXCL16^+^ cells led to a significant increase of proliferative T cells which have divided two times (43.6% of CD4 and 69% of CD8) when compared to T cells co-cultured with activation beads (17% of CD4 and 21.5% of CD8). However, most of the T cells co-cultured with the activation beads (57.5% of CD4 and 66% of CD8) divided more than two times whereas division of the T cells co-cultured with CXCL16^+^ cells was halted in the second division phase (Fig. S4C).

Cxcl16 has been reported to be a chemokine attracting immune cells such as T cells (Agostini et al., 2005; Matloubian et al., 2000), macrophages (Kim et al., 2019), neutrophils (Woehrl et al., 2010), and monocytes (Allaoui et al., 2016) in various disease contexts. To examine the potential role of myeloid-derived *Cxcl16* in regulating tumor infiltration of bone marrow-derived immune cells (BMD cells), we performed an intratumoral *Cxcl16* neutralization experiment (Fig. 4C and Methods). In this experiment, after treating the mice either with Vehicle or MCT for two weeks, the mice were intratumorally (i.t.) injected with three doses of either IgG or CXCL16 neutralizing antibody (NAb). On the second day of antibody injection, BMD cells were adoptively transferred into the tumor bearing mice (Fig. 4C). CXCL16 NAb injection significantly decreased the overall percentage of tumor-infiltrated CXCL16^+^ CD45.1^+^ cells in MCT-treated tumors (Fig. 4D and Fig. S4D). Next, to explore the impact of CXCL16 NAb on lineage dynamics of tumor infiltrating lymphocytes (TILs), we looked into percentages of CD45.1^+^ progenitor-like c-KIT^+^LIN^-^, Cluster 0-like CD11b^high^PD-L1^high^, and CD11c^high^ I-A-I-E^high^ DCs (Fig. S4E, gating strategy). CXCL16 NAb injection did not significantly alter the infiltration of CD11b^high^PD-L1^high^ cells in PBS tumors while the decreasing trend was more apparent in the MCT-treated tumors (Fig. 4E, left). In contrast, Cxcl16 NAb i.t. injection significantly decreased the infiltration of c-KIT^+^LIN^-^ CD45.1^+^ cells in both PBS and MCT-treated tumors (Fig. 4E, center). More interestingly, the infiltration of CD11c^+^ I-A-I-E^high^ DC into the MCT-treated tumors (Fig. 4E, right) but not in PBS-treated, also significantly reduced after Cxcl16 NAb i.t. injection. Above results suggest the CXCL16 in the TME has a more prominent role in attracting progenitor-like c-KIT+LIN-myeloid cells as well as peripheral DCs into the MCT-treated tumors.

CXCL16^+^ DCs are known to interact with T cells, especially CXCR6^+^ T cells (Di Pilato et al., 2021). Hence, we examined the T cell infiltration into the tumors upon CXCL16 neutralization (Fig. 4SF, T cell gating). CXCL16 NAb did not alter the infiltration of CD4^+^ T cells in the PBS-treated tumors but increased in MCT-treated tumors (Fig. 4F, left). Among these CD4^+^ T cells, CXCL16 NAb significantly decreased PD-1^high^ CD4^+^ T cells in MCT-treated tumors (41% in IgG to 11% in NAb) (Fig. 4F, middle-left). In addition, CXCL16 NAb more significantly reduced the CD8a^+^ T cells infiltration in the MCT-treated tumors (44.6% in IgG and 12% in NAb group), but not PBS-treated tumors (Fig. 4F, middle-right). Similarly, among these CD8a^+^ T cells, CXCL16 NAb reduced the PD-1^high^ CD8^+^ T cell infiltration specially in MCT-treated tumors (57% in IgG to 20.5% in NAb group) (Fig. 4F, right). CXCL16 NAb did not significantly alter the infiltration pattern of CD4^+^ CXCR6^high^ or CD8^+^ CXCR6^high^ cells in MCT-treated tumors (Fig. S4G, box plot). Collectively, our results suggest that the increase of CXCL16, mostly derived from Path 4 clusters cells, attracts exhausted PD-1^high^ T cells into the MCT-treated tumors.

Previous studies showed CXCL16 blockage has therapeutic potential in curbing thyroid tumor growth (Kim et al., 2019). To investigate the therapeutic potential of combining immunotherapy and CXCL16 blockage, we combined PD-1 checkpoint inhibitor with CXCL16 NAb (Fig. 4G, Treatment scheme). The C3-1-TAg TNBC primary tumor bearing mice were chemoprimed with MCT for two weeks and then treated with anti-PD-1 antibody in combination with either IgG or CXCL16 NAb. The mean percentages of tumor-associated CXCL16^+^ CD45^+^ cells among all the CD45^+^ cells decreased from 8.5% in IgG to 2.1% in CXCL16 NAb group (Fig. 4H). Intriguingly, the mean relative tumor volume significantly decreased from 4 in IgG to 1.8 with combo treatment of CXCL16 NAb and anti-PD-1 antibody compared with anti-PD-1 alone (Fig. 4I). Next, we examined TIME dynamics under CXCL16 NAb treatment, including CD4^+^ and CD8^+^ T cells, CD11c^high^ I-A-I-E^high^ DCs, CD11b^high^PD-L1^high^, and Cluster 0 cells (Fig. S4H and S4I, gating strategy). Both CD4^high^ T (16.4 to 40.5%) and CD8^high^ (5 to 17%) T cells significantly expanded upon combo treatment compared with anti-PD-1 single treatment (Fig. 4J) partially explains the reduced tumor size. While the increase of tumor infiltrating T cells is encouraging, upon CXCL16 NAb treatment, the CD11c^+^ I-A-I-E^high^ DCs decreased (10% in IgG to 3% in CXCL16 NAb). While the percentage of CD11b^high^PD-L1^high^ cells among CD45+ cells did not change, treatment of CXCL16 NAb led to an increase of Cluster 0 cells under MCT plus anti-PD-1 dual treatment (Fig. 4K), suggesting a shift in balance of the trajectory towards Path 1 Cluster 0 (*Cd274+, Cd14+, S1008/9+*) compensatory to the inhibition of CXCL16 signaling.

### STAT1 regulates PD-L1 expression in Cluster 0 cells and influences T cell activation status

The shift towards Path 1 after CXCL16 NAb treatment suggests Path 1 Cluster 0 cells might confer resistance to the anti-PD-1 ICB treatment. To examine whether Cluster 0 cells could suppress the T cell activation and expansion on T cell activity, we performed an in *vitro* co-culture assay where we co-cultured the MCT-tumor infiltrating Cluster 0 cells with T cells in combination with T cell activation beads (Fig. 5A and Fig. S5A). To enrich the Cluster 0 cells from tumor infiltrating immune cells, we first examined the top 20 differentially up-regulated genes (**Supplementary Table 3**, Cluster 0 marker tab). Among Cluster 0 marker genes, we chose to use cell surface protein CCR1 for antibody-mediated cell enrichment as it shows a more specific expression in Cluster 0 than CD14 (Fig. S3A). We magnetically enriched CD45^+^ TIL using Mojosort^TM^ and further used CD11b and CCR1 as markers to sort out CD11b^+^CCR1^+^ Cluster 0 cells using FACS (Methods). Of all the CD45^+^ TIL cells, about 10.7% are CD11b^+^CCR1 high (Fig. 5A, biaxial plot). 81% of naive CD4^+^ T cells proliferated more than once when co-cultured with T cell activation beads and IL2 (Fig. 5B, box plot, gating shown in Fig. S5A). In the presence of CCR1^high^ myeloid cells, the % of proliferated CD4^+^ T cells did not change (mean of 72%). However, in the presence of CCR1^low^ myeloid cells in the co-culture, the % of proliferated CD4^+^ T cells significantly decreased (mean of 18%). Similarly, 70.8% of naive CD8^+^ T cells proliferated more than once when co-cultured with T cell activation beads and IL2. While the presence of CCR1^high^ myeloid cells did not change (mean of 70%) their proliferation, CCR1^low^ myeloid cells decreased the naive CD8^+^ T cell proliferation significantly (mean of 20%) (Fig. 5B, right box plot). This observation suggested that CCR1^high^ (Cluster 0) myeloid cells do not have MDSC-like properties which inhibit T cell proliferation. Instead, co-culture with CCR1^high^ myeloid cells led to a significant increase in the ratio of proliferating to non-proliferating PD-1^high^ CD4^+^ or CD8^+^ T cells (Fig. 5C). Importantly, this observation was specific to T cells co-cultured with Ccr1^high^ myeloid cells but not Ccr1^low^ myeloid cells (Fig. 5C), suggesting Cluster 0 cell pushes T cell toward exhaustion-like status. To further comprehend the transcriptome of T cells, we identified CITE-CD45 and CITE-CD3^+^ TIL cells from our CITE-seq data and clustered them into 10 Seurat Clusters, of which 1 and 2 showed a high expression of *Cd3e*. Then these cells were further subsetted and reclustered, into three Seurat Clusters including *Cd8a*^+^ T cells (Cluster 0), *Cd4*^+^ T cells (Cluster 1), and *Foxp3*^+^*Cd4*^+^ T cells (Cluster 2) (Fig. S5B, left). Upon MCT-treatment, some T cell dysfunctional markers such as *Ikzf4*, *Ctla4*, and *Lag3* increased in both *Cd8a* and *Foxp3 Cd4* T cell clusters, *Klrg1* and *Tox* increased in *Foxp3 Cd4*, and *Pdcd1* increased in *Cd8a* T cell cluster (van der Leun et al., 2020), suggesting a transcriptional shift toward exhaustion in vivo.

**Figure 5:**
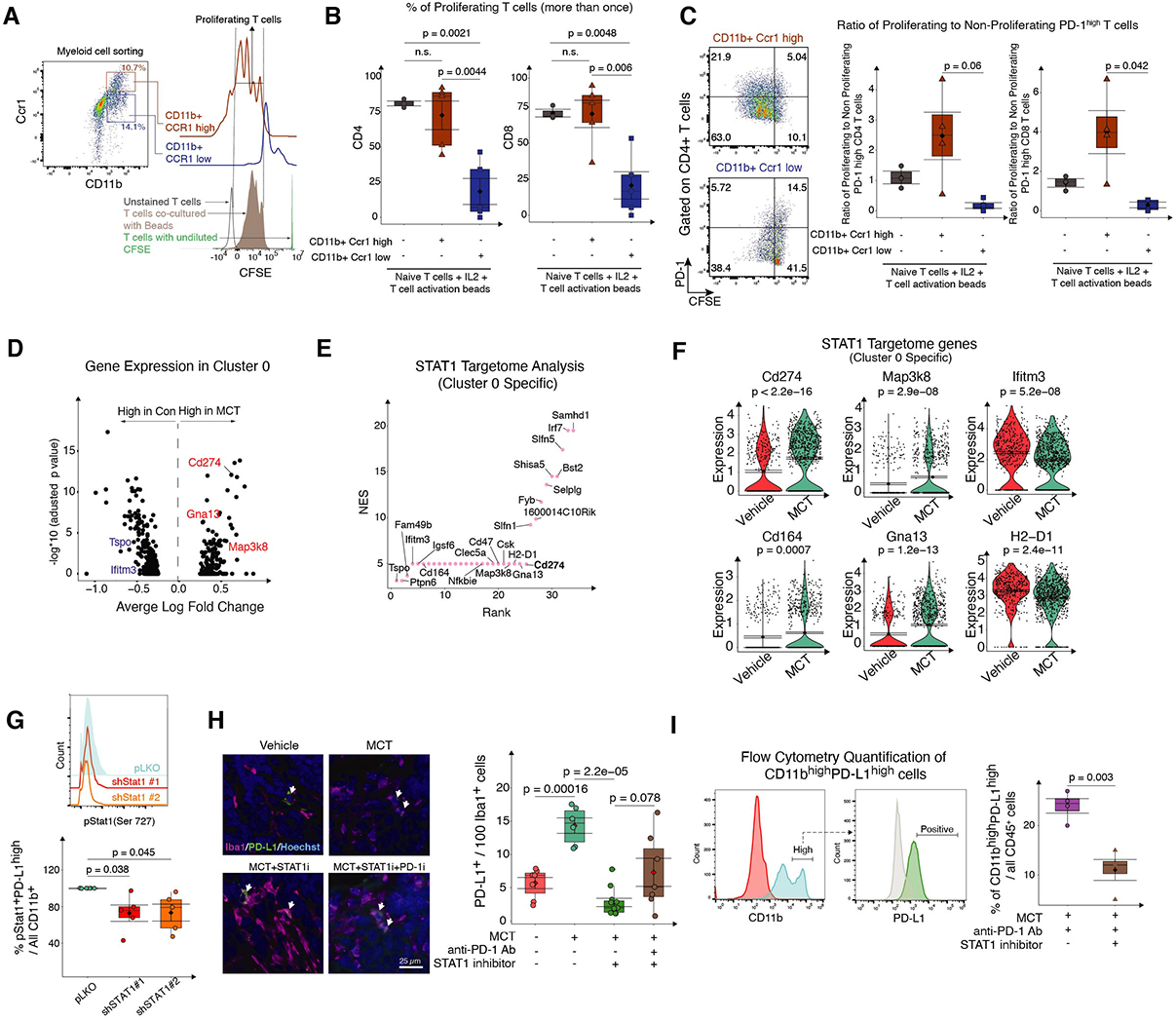
STAT1 regulates PD-L1 expression in Cluster 0 cells and influences T cell activation status. (A) Gating strategy to identify FACS sorted CD11bhigh CCR1high, CD11bhigh CCR1low myeloid cells and CFSE levels on CD8+ T cells in co-culture. (B) Box plots show % of proliferating CD4 and CD8 T cells in coculture with T cell activation beads with or without myeloid cells. (C) Biaxial plots (left) show the gating strategy to identify the PD-1 levels on CFSE diluted and proliferating T cells. Box plots (right) show the ratio of PD-1high proliferating and non-proliferating CD4 and CD8 T cells. (D) Volcano plot shows DGE on Cluster 0 cells when MCT-treated. (E) Scatter plot shows the potential gene targets (NES > 3 and p < 0.05 DEG between MCT and Vehicle treated) in Cluster 0 specific STAT1 targetome. (F) Violin plots show expression of Cd274, Map3k8, Ifitm3, Cd164, Gna13, and H2-D1 genes by Cluster 0 cells (each dot is a cell and cells are from n = 3 biological repeats pooled together). (G) Histogram shows the levels of pStat1 in CD11b+ cells in different transfection conditions. Box plot shows % of pStat1+ PD-L1+ CD11b+ cells after shRNA STAT1 transfection relative to pLKO transfection (n = 5 biological repeats). (H) Representative IF images of Iba1 and PD-L1 stained tumors treated with either Vehicle, MCT, MCT + STAT1 inhibitor, or MCT + STAT1 inhibitor + PD-1 inhibitor (left). Box plot (right) shows the quantified Iba1+ PD-L1+ cells of all the Iba1+ cells (n ≥ 3 biological repeats). (I) Histograms show flow cytometry gating strategy and box plots show quantification of PD-L1+ CD11b+ cells of all the CD45+ immune cells in dual-therapy (MCT + PD-1 inhibitor) and tri-therapy (MCT + PD-1 inhibitor + STAT1 inhibitor) treated TNBC tumors (n = 4 biological repeats).

We next analyzed Cluster 0 specific DGE (differential gene expression), DGEP (differential gene expression pathway), and *Stat1* targetome. Upon MCT treatment, Cluster 0 cells up-regulated marker genes related to innate immunity (*Gna13*) and immune suppression (*Cd274*) (Sharma and Allison, 2015) (Fig. 5D and Fig. S5C). Furthermore, GSEA analysis of Cluster 0 specific DGE correlated with a downregulation of adaptive immune system pathways, including CD8^+^ T cell activity, and dendritic cell maturation (Fig. S5D). Cluster 0 cells also up-regulated expression of *GPCR* ligand binding and *GPCR* activation pathways that are critical for immune cell functionality (Tischner et al., 2017) and *TGFβ* pathway, known to create an immune suppressive TIME (Batlle and Massagué, 2019) (Fig. S5D). Since *STAT1* is the major transcription factor that drives Cluster 0 transcriptome (Fig. 3E, SCENIC analysis), we further examined potential gene targets of *STAT1* (Targetome) that are differentially expressed by Cluster 0 (Fig. 5E). SCENIC based gene regulatory network analysis (GRN) (Aibar et al., 2017) correlated more than 200 genes (NES > 3) under *STAT1* targetome (**Supplementary Table 8**). Interestingly, in addition to *Cd274*, Cluster 0 specific *STAT1* targetome also contains genes that are immune-related, including *Tspo*, *Ifitm3*, *Cd164*, *Clec5a*, *H2-D1*, *Gna13*, and *Irf7* (Fig. 5E). MCT significantly increased the expression of a T cell co-inhibitory signal, *Cd274* (Sharma and Allison, 2015), Map kinase protein *Map3k8,* metastasis promoter, *Cd164* (Samanta et al., 2012), and *Gna13* G protein (Rasheed et al., 2018) (Fig. 5F). Expression of interferon-regulatory genes *Ifitm3* (Kim et al., 2020), histocompatibility antigen *H2-D1*, innate immune-related *Clec5a*, immunomodulatory genes *Cd47*, *Tspo*, and *Irf7* decreased with MCT treatment (Fig. 5F and S5E). Next, we validated *STAT1’s* role in regulating tumor-associated PD-L1^+^ myeloid cells by knocking down *STAT1* in tumor-associated CD11b^+^ cells. Tumor bearing C3-1-TAg mice were treated with MCT for 28 days. Tumor-associated CD11b^+^ cells were then transfected with lentivirus carrying either pLKO shRNA control or *STAT1* targeting shRNAs (**Methods**). Compared with the pLKO control shRNA group, shRNA *STAT1* led to a decrease in percentage of CD11b^+^ cells with high phospho-Stat1 (pStat1) levels (Fig. 5G and Fig. S5F, gating strategy). Furthermore, multigene association analysis of TCGA and GTex projects’ RNA-seq data showed a stronger association between *STAT1* and *Cd274* expression in the invasive breast carcinoma (R = 0.72) compared with the correlative pattern in healthy mammary gland tissue (R = 0.3). Similarly, the association between *STAT1* and *Cd274* was stronger in cancerous tissues in multiple cancer types (Fig. S5G).

Given the close association between PD-L1 levels and *STAT1* (phosphorylation) levels in Cluster 0 cells, we treated mice with a combination of *STAT1* inhibitor Fludarabine (Frank et al., 1999) with or without PD-1 inhibitor along with MCT. PBS-treated tumors served as a control group for MCT-treated tumors. IF results suggested that out of every 100 Iba1^+^ cells the frequency of PD-L1^+^ Iba1^+^ cells increased from 5% in Vehicle-treated tumors to 14% in MCT. Furthermore, combining *STAT1* inhibitor with MCT decreased their frequency to 3% and combining *STAT1* and PD-1 inhibitors with MCT decreased the frequency to 10% (Fig. 5H, box plot). Flow cytometry results suggested that among all the CD45^+^ cells, the mean frequency of PD-L1^+^ CD11b^+^ cells significantly decreased from 24% to 11% when *STAT1* inhibitor was added to the regimen (Fig. 5I).

### Modulating STAT1 signaling enhances anti-PD-1 antibody mediated anti-tumor responses

To examine the impact of PD-1 inhibitor on tumor growth, we treated MMTV-neu and C3-1-TAg spontaneous tumor bearing mice with PD-1 immune checkpoint blockade (ICB) after two weeks of MCT treatment (Fig. 6A, Treatment Regimen 1). While MCT treatment significantly reduced tumor size, the subsequent anti-PD-1 ICB treatment alone was not sufficient to further enhance the antitumor efficacy of MCT in both tumor models (Fig. 6B). We reasoned that, based on our understanding of the TME immune cell heterogeneity (Fig. 2-5), targeted modulation of TME immune components after MCT treatment could potentially further potentiate the ICB treatment.

**Figure 6.**
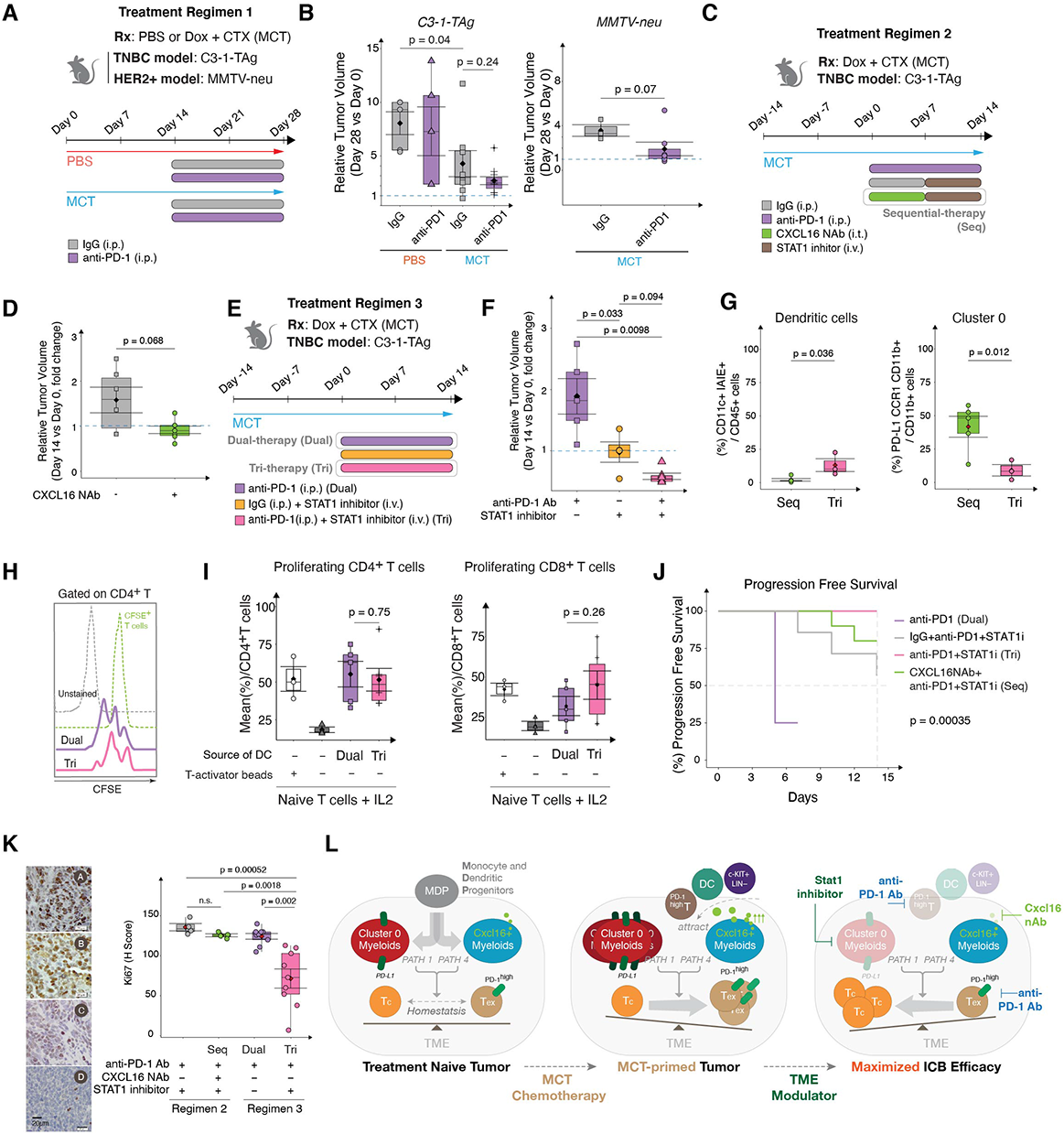
Modulating STAT1 signaling enhances anti-PD-1 antibody mediated anti-tumor responses. (A) Treatment scheme for synergizing anti-PD-1 with MCT (Regimen 1). (B) Box plots show tumor volumes on Day 28 of treatment (Regimen 1) relative to Day 0 in C3-1-TAg (left) and MMTV-neu (right) mice. 3 to 5 mice pooled over three cohorts (n ≥ 3 per group). (C) Treatment scheme for synergizing CXCL16 NAb with STAT1i response in MCT and anti-PD-1 treated mice (Regimen 2). (D) Box plots show tumor volumes on Day 14 of treatment (Regimen 2) relative to Day 0 in C3-1-TAg mice. 3 to 5 mice pooled over three cohorts (n ≥ 5 per group). (E) Treatment scheme for synergizing STAT1i and anti-PD-1 antibody response (Regimen 3). (F) Box plots show tumor volumes on Day 14 of treatment (Regimen 3) relative to Day 0 in C3-1-TAg mice. 3 to 5 mice pooled over three cohorts (n ≥ 5 per group). (G) Box plots quantify flow cytometry identified dendritic cells and Cluster 0 cells in Seq-therapy and Tri-therapy. (H) Flow cytometry derived gating strategy identifies CFSE levels on CD4+ T cells. (I) Box plots show % of CD4+ (left) and CD8a+ (right) T cells proliferating in co-culture with tumor-associated cDC2s (CD11b+CD11c+ cells) isolated after treatment with either Dual or Tri-therapy (n = 3 to 5 biological repeats). (J) Progression free survival probability in different treatment regimens showing time taken for the tumor volume to increase from Day 0 of treatment to Day 14. (K) Representative IHC images of Ki-67 stained TNBC tissues treated with either Regimen 2 or 3 (left). Box plot (right) quantifies H-Score (n ≥ 3 biological replicates per group). Data in box plots represent n ≥ 3, Mean ± SEM, Diamonds Means, Horizontal black lines Medians, and T-test p-values. Survival pot was generated using the Survimer package in R. (L) Schematic shows the proposed-model of MCT-mediated immune modulation in breast tumor.

We have previously observed 1) intratumoral CXCL16 neutralization (CXCL16 NAb) improved the efficacy upon MCT plus anti-PD1 treatment (Fig. 4I); 2) CXCL16 neutralization expanded tumor-associated Cluster 0 cells while decreasing the percentage of overall dendritic cells (Fig. 4K); 3) STAT1 inhibition led to a decrease of PD-L1 expression on myeloid cells (Fig. 5G-I). Thus, we designed a sequential treatment regimen of CXCL16 NAb and STAT1 inhibitor to test possible enhanced efficacy of sensitizing MCT-primed tumor to anti-PD-1 treatment. The C3-1-TAg mice bearing primary breast tumors were MCT-primed for two weeks and then treated with anti-PD-1 antibody in combination with either IgG or CXCL16 NAb prior to STAT1 inhibitor treatment (Fig. 6C, Treatment Regimen 2, **Seq**). The mean percentages of tumor-associated CXCL16^+^ Iba1^+^ myeloid cells among all the Iba1+ myeloid cells decreased from 17.4% in IgG group to 2.7% in CXCL16 NAb group (Fig. S6A). However, one week treatment of CXCL16 NAb prior to STAT1 inhibitor (Treatment Regimen 2) was only able to moderately improve the efficacy of Tri-therapy and maintain a stable disease over the course of treatment (Fig. 6D). The potential increase of Cluster 0 cells after CXCL16 NAb might counteract the overall anti-cancer response. Thus, we further hypothesized that targeting STAT1 right after MCT-priming could potentially improve anti-PD1 treatment efficacy (Fig. 6E, Treatment Regimen 3). Compared with the baseline efficacy (MCT + anti-PD1), adding STAT1 inhibitor (Tri-therapy, Tri) significantly improved anti-tumor efficacy of anti-PD1 treatment (Dual-therapy, Dual) (Fig. 6F, p = 0.0098). Given such significantly improved efficacy (Fig. 6F), we sought to examine the possibility that STAT1 inhibition in MCT-primed tumors modulates tumor-associated myeloid cells. Intriguingly, the tumor-associated dendritic cell percentage (Fig. 6G, Fig.S6B) increased and Cluster 0 cells (CD11b+ PD-L1+ CCR1+, Fig. 6G, Fig.S6B) percentage decreased in Tri-therapy (Tri) used in regimen 3 when compared to Sequential (Seq) treatment used in regimen 2. This suggests the potential shift of myeloid cell trajectory Path 1 (Cluster 0 dominated) to Path 4 (Dendritic cell dominated) when applying the Tri-therapy (Tri) scheme.

To examine the influence of dual or tri-therapy on cDC2 interaction with T cells, we performed *in vitro* co-culture based T cell proliferation assay (Fig. 6H-I). cDC2-like (Cd11c^+^Cd11b^+^) cells were isolated from tumors that have been treated with either dual or tri-therapy, then co-cultured with naive T cells. Co-culture with cDC2s isolated either from tri-therapy treated tumors did not increase CD4^+^ T cell proliferation when compared to dual therapy (Fig. 6I, left). cDC2s isolated from dual or tri-therapy treated tumors did not sufficiently support CD8a^+^ T cell proliferation *in vitro,* despite a trend of increasing proliferation in the tri-therapy group (45%) (Fig. 6I, right). Above observation suggests that the improved anti-tumor efficacy of the STAT1 inhibitor-containing Tri-therapy might primarily be due to a reduced Cluster 0 cells, which drives the excessive T cell exhaustion (Fig. 5C), rather than increased cDC2 number and activity. Overall, the tri-therapy improved the progression-free survival when compared to other strategies in either of the regimens (Fig. 6J, Pink line). The H-score of the Ki-67 nuclear stain decreased most significantly from 120 in dual-therapy to 72 in tri-therapy (Regimen 3) treated tumors (Fig. 6K).

## DISCUSSION

### A trajectory view of MCT-primed TIME in potentiating ICB immunotherapy

MCT regimen was originally conceived as an alternative chemotherapy regimen to MTD with potential anti-angiogenic benefits (André et al., 2014). In addition to the expected anti-angiogenic effect, MCT stimulates multiple antitumorigenic immune cell subsets in the context of breast cancer (Munzone and Colleoni, 2015). Low-dose chemotherapy has been demonstrated to have immune stimulatory properties in the TONIC metastatic TNBC clinical trial and improved overall response rate in 20% (35% of Doxorubicin treated) of participants to PD-1/PD-L1 blockade (Voorwerk et al., 2019). Despite a generally low response rate of breast cancer to ICB, this study highlighted the clinical significance of using chemotherapy as a “primer” for immune checkpoint blockades even in the late-stage metastatic setting. However, while the improvement seen in the TONIC study is encouraging, overall anti-tumor efficacy remains limited. A large portion of patients were still not responsive to anti-PD1 ICB treatment (Voorwerk et al., 2019). In our preclinical setting, addition of anti-PD-1 immunotherapy to MCT regimen was only marginally improved anti-tumor efficacy (Fig. 6A-B, regimen 1), suggesting other potential immune suppressants within the TIME that restrict the full potential of ICB’s efficacy. To reveal this dynamic equilibrium of TIME after MCT, we used CITE-seq and trajectory analysis to map the single-cell immune landscape of breast tumor after low-dose MCT-priming (Fig. 2). It is increasingly appreciated, besides tumor-T cell direct engagement through immune checkpointmolecules (Wei et al., 2018), that the maximal benefits of ICB treatment requires T cell-dendritic cell crosstalk (a “licensing” model) (Garris et al., 2018). Tumor-infiltration by DCs also correlates with immunotherapeutic efficacy (Engblom et al., 2016; Garris et al., 2018). T-DC crosstalk sensitizes the tumor to ICBs (Garris et al., 2018) and ensures durable responses in melanoma and lung cancer (Garris et al., 2018; Hargadon et al., 2018). Our single cell analysis showed that MCT enhances T and myeloid cell colocalization (Fig. 1C-E), expands antigen presenting cDCs (Fig. 1G-H). However, MCT-priming also parallely enables the evolution of a subset of paradoxical myeloid cells (Cluster 0 cells), expressing both immune-stimulating and immune-suppressive transcriptional profile, leading to accelerated T cell exhaustion (Fig. 2B, 3A, S3A-H, 5D-E). In contrast to an over-simplified binary models of either immune stimulating or suppressive tumor microenvironment, this coexistence of antigen presenting and immunosuppressive myeloid cells underscores the notion of *de novo* immune homeostasis achieved in MCT-primed TIME, highlighting a resilient nature of tumor immune microenvironment to re-establish homeostasis in response to the systemic stimulation - a tenet in biology at the organismal level (Yosef and Regev, 2016).

### Intratumoral CXCL16 inhibition in MCT-primed TIME potentiates anti-tumoral effects of ICB

CXCL16 has been shown to improve tissue tolerance, and recruit T cells in various physiological and pathological contexts (Agostini et al., 2005; Jiang et al., 2005; Wang et al., 2018; Woehrl et al., 2010). In ovarian, thyroid, and lung cancers CXCL16 promotes tumor progression (Kim et al., 2019; Srivastava et al., 2021; Veinotte et al., 2016). Immunogenic chemotherapy promoted CAR-T cell recruitment by increased CXCL16 secretion in lung cancer (Srivastava et al., 2021). Furthermore, CXCL16-CXCR6 interaction promoted tolerance to cardiac allograft transplants and successful fetal development (Jiang et al., 2005; Wang et al., 2018). In our study, we observed that MCT increased CXCL16 in the dendritic cell specific trajectory paths (Fig. 4A,B and Fig. S4B). Neutralization of CXCL16, resulted in decreased tumor-infiltration of immune cells, specifically CD45^+^ CD11b^+^ PD-L1^high^ cells, c-Kit^+^Lin^-^ progenitor cells, PD-1^high^ T cells (Fig. 4D-F). Reducing the recruitment of peripheral PD1^high^ and PD-L1^high^ cells to TIME partially explains the improvement of the efficacy of anti-PD-1 antibody upon intratumoral neutralization of CXCL16 (Fig. 4I-J). However, the tumor continues to progress (Fig. 4I, Relative Tumor Burden >1.5 fold increase, Day 28 vs Day 14), potentially due to an significant increase of Cluster 0 cells after CXCL16 neutralization (Fig. 4K), suggesting the dynamic nature of the TIME in response to immune modulator CXCL16. Therefore, in order to maximize the ICB’s anti-cancer efficacy, a holistic and dynamic view of TIME is needed.

### Targeting STAT1-PD-L1 axis on tumor-associated myeloid cells to shift the balance between immune stimulator and suppressor pathways

Immune cell plasticity drives both anti- and pro-tumor immunity to maintain the TIME homeostasis. Immune plasticity, exemplified in the highly plastic myeloid cells, could be leveraged to improve the efficacy of anticancer therapies (Yosef and Regev, 2016). Through tumor-associated myeloid cell subset trajectory inference and SCENIC regulon analyses, our study revealed the emergence of immune deterring myeloid cluster 0 after MCT, which is transcriptionally driven by *STAT1* regulon. Interestingly, it has been shown that eIF4F-regulated translation of *STAT1* mRNA is a critical step of IFN-γ-induced PD-L1 expression on melanoma tumor cells, partially contributing to tumor immune evasion mechanisms (Cerezo et al., 2018), suggesting a translational significance of modulating elF4F-STAT1 pathway as a potential cancer immunotherapeutic modality in melanoma patients (Cerezo et al., 2018). Complementary to *Cerezo et al.* study, our study further demonstrated *STAT1* signaling as a dominant regulon driving the tumor-infiltrated immature myeloid cells (Cluster 0) after MCT-priming (Fig. 3C, 3E). The role of *STAT1* signaling in TME has been previously suggested in Renca monoclonal cancer model, where a high expression of inflammatory gene signature driven by *STAT1* signaling in TME is correlated with the tumors’ responsiveness to ICBs (Zemek et al., 2019). In the MCT-primed breast cancer context, we observed similar inflammatory signatures in TME cells (Fig. 3A, S3A, 5D-F). However, despite an enriched inflammatory signature in the TME, MCT-priming only marginally improved the response of breast cancer to anti-PD-1 treatment (Fig. 6B). Our single-cell trajectory analysis shed the light on the root cause of this suboptimal outcome and demonstrated an intrinsic dual-role of chemotherapy priming - inducing a potentially immunogenic inflammatory signature while concurrently stimulating *STAT1* regulon in TME to facilitate local immune tolerance through up-regulating immunosuppressants due to CXCL16^high^ dendritic cells (Fig. 4) and Cluster 0 immature myeloid cells (Fig. 5). Such MCT-induced tolerance imposed by CXCL16^high^ dendritic cells and Cluster 0 myeloid cells partially explains the only subtle improvement of therapeutic efficacy seen on combining MCT with anti-PD-1(Fig. 6B), highlighting the nature of local immune equilibrium regulated by collective factors, rather than an over-simplified PD-1/L1 centric binary model of immune regulation. Therefore, targeting *STAT1* - a master regulator of collective immunosuppressants - in myeloid Cluster 0 cells using *STAT1* inhibitor in MCT-primed breast tumor effectively shifted the immune homeostasis in the favor of immunogenicity, which resulted in a much-improved response to ICB (Fig. 6F, J, L).

In summary, enabled by multimodal single-cell analysis, we uncovered the pro-tumoral immune suppressive myeloid cells within the MCT-primed tumor microenvironment. Our study provided mechanistic insights and preclinical rationale for targeting the *STAT1* regulon as a neoadjuvant regimen for MCT-primed breast tumors to achieve the maximum clinical benefit of the anti-PD1 immunotherapy.

## MATERIALS AND METHODS

### Mice

To model the clinical scenario, we used spontaneous tumor bearing immunocompetent mouse strains, MMTV-neu, MMTV-PyMT (Guy et al., 1992), and C3-1-TAg (Maroulakou et al., 1994), representing HER2 overexpressing and TNBC in clinic respectively. For MMTV-neu, FVB/N-Tg(MMTVneu)202Mul/J (Jax Stock, 002376) homozygous males were crossed with homozygous females. For spontaneous tumor developing MMTV-PyMT B6 mice, FVB/N-Tg(MMTV-PyVT)634Mul/J carrying MMTV-LTR driving the mammary gland specific Polyoma middle T antigen is backcrossed to C57BL/6J (Jax Stock, 022974). For spontaneous TNBC tumor development, C3-1-TAg males hemizygous for Tg(C3-1-TAg) (Jax Stock, 013591) were crossed with noncarrier (FVB) females. For non-tumor bearing C3-1-TAg mice (Aprelikova et al., 2016), C3(1)/Tag-REAR (Jax Stock, 030386) mice (Hemizygous crossed with noncarrier) were used as tumor transplantation recipients as well as used for peripheral blood analyses. All mice were bought from the Jackson Laboratory and bred at Freiman Life Sciences Center, University of Notre Dame. Female mice were used for all the experiments. All the experiments were performed as per University of Notre Dame’s Institutional Animal Care and Use Committee (IACUC) approved protocols.

### Drugs and treatment dosages

Doxorubicin (DOX, MedChem Express, HY-15142) and Cyclophosphamide (CTX, MedChem Express, HY-17420) were used in the chemotherapy cocktail. The clinically used doses of DOX and CTX were converted from human to mouse equivalents using FDA-approved formula, HED (mg/kg) = Animal Dose (mg/kg) * 0.08. Chemotherapy cocktail was administered intravenously either once a week (MTD) or thrice a week (once every two and a half days) (MCT) for a period of four weeks. The dosage of DOX used was 2 mg/kg body weight and CTX 20 mg/kg body weight for the MTD cohort and a third of the MTD dose for the MCT cohort. For a PD-1 inhibitor treatment trial, both PD-1 inhibitor (BioXcell, BE0146) and IgG (BioXcell, BE0087) antibodies were injected intraperitoneally once a week at a dose of 150 µg per mouse. The PD-1 inhibitor was dosed at 6 mg/kg body weight, which is a mouse equivalent dose of PD-1 inhibitor clinical human dose. *Stat1* inhibitor (MedChem Express, HY-B0028) was given as 8.4 mg/kg body weight (derived from clinically used human doses and calculated as per mouse equivalents) in MCT strategy intravenously along with MCT as a cocktail prepared right before administration.

### Single-cell isolation for CyTOF and CITE-seq analysis

Commercially available Collagenase Type 1 enzyme powder (Thermo Fisher Scientific, 17100-017) dissolved in DMEM High Glucose Medium (1% Penicillin/Streptomycin and 10% Fetal Bovine Serum) (Thermo Fisher Scientific, 11965092) was used to digest the tumors. The tumors were enzymatically digested for 60 min after mincing. Live cells were enriched using Ficoll solution (GE Healthcare, 17-5446-02). For density gradient based enrichment of cells, 4 mL of cell suspension was added gently as a layer on the top of 3mL of Ficoll Paque solution in a 15 mL conical tube. The conical tube was centrifuged at 500 RCF with the brake off at 18 ℃. All the cell layers were collected. The pellets with blood cells and granulocytes were subjected to red blood cell lysis and granulocytes separated. Red blood cells (RBCs) were lysed using the ACK Lysis buffer (Lonza, 10-548E). Dead cell removal kit (Miltenyi Biotec, 130-090-101) was used to remove dead cells. CD45 magnetic beads (Miltenyi Biotec, 130-052-301) and or CD11b magnetic beads (BioLegend, 480109) were used to isolate the CD45^+^ and CD11b^+^ cells. For isolating splenocytes, the spleen was mashed in heat inactivated RPMI medium. RBCs were lysed in 1X red blood cell lysis buffer ACK Lysis Buffer (Lonza 10-548E). Either CD45 magnetic beads or T cell isolation kit (Miltenyi Biotec, 130-095-130 or BioLegend, 480024) was used to isolate cells of interest. Cells were incubated at 4℃ to maintain their viability and passed through a 40 µm cell strainer (VWR, 76327-098) to dissociate single cells from clumps. Single cells were maintained viable and separate in buffers with 5% Bovine Serum Albumin (BSA) and 2mM Ethylene Diamine Tetra Acetic Acid (EDTA).

### Blood collection for peripheral immune cell quantification

Whole blood was collected from an anesthetized mouse (100 µL) into an anticoagulant coated syringe. RBCs were lysed using ACK Lysis Buffer (Lonza 10-548E) and the blood was further diluted at 1:100 ratio in the cell staining buffer (Biolegend, 420201). The cells were stained with CD45 flow antibody and proceeded for flow cytometry to analyze the frequency of CD45^+^ immune cells.

### Flow cytometry and fluorescence assisted cell sorting (FACS)

Following single-cell isolation, cells were washed in the cell staining buffer (Biolegend, 420201). For experiments on cell surface antibody staining, F_C_ and F_ab_ receptors were blocked through incubation with 100 uL cell staining buffer for 30 min at 4°C and then stained with antibodies of interest for 15 min at 4°C. For intra-cytoplasmic staining, the cells were blocked for non-specific binding using the cell staining buffer and then were fixed and permeabilized using True Phos Perm Buffer (BioLegend, 425401) before antibody staining. The antibodies used for analysis on FC500 were: PE/APC anti-mouse CD45 antibody (BioLegend, 103105/103112), Alexa Fluor 488 anti-mouse CD11c antibody (BioLegend, 149021), APC/Cy7 CD11b antibody (BioLegend, 101225), APC anti-mouse CD103 (BioLegend, 121413), APC/Cy7 CD8 anti-mouse antibody (BioLegend, 100713), PE anti-mouse PD-L1 (BioLegend, 124308), Alexa Fluor 488 pStat1 (Ser 727) (Biolegend, 686410), PE anti-CD44 anti-mouse antibody (BioLegend, 10323). For analysis on Cytek Flow cytometer, Myeloid cell specific markers used were: BV421 anti-mouse CD86 (BioLegend,105031), BV570 anti-mouse CD11c (BioLegend,117331) or 11b (BioLegend,101233), BV605 anti-mouse CD11b (BioLegend,101257) or 11c (BioLegend,117333), BV711 anti-mouse PD-L1(BioLegend,124319), FITC anti-mouse Ccr1 (BioLegend,152505), PE anti-mouse CXCL16 (BD, 566740), PE-Dazzle/594 I-A-I-E (BioLegend,107647), PerCP-Cy5.5 anit-mouse pStat1(p727) (BioLegend,686415), PE-Cy7 CD45 (BioLegend,157206); Lineage specific markers used were Pacific Blue anti-mouse Lin (BioLegend,133305) and BV510 anti-mouse c-KIT (BioLegend,105839); T cell specific markers used were BV 421 PD-1 (BioLegend,135217), BV510 CD69 (BioLegend,104531), FITC anti-mouse CD45 (BioLegend,103122), PE/Cy5 CD4 (BioLegend,100410), PE-Dazzle 594 CD8a (BioLegend,100762), PerCP-Cy5.5 CXCR6 (BioLegend,151120), PE-Cy7 anti-mouse CD3e (BioLegend,100319) and immune cell marker BV785 CD45.1 (BioLegend,110743). Following staining, cells were washed and resuspended in a 250-300 µL cell staining buffer. The cells were sorted on either a BD Biosciences 1108 FACS Aria III sorter or analyzed on BD FC500 flow cytometer or Cytek Northern Lights (Cytek NL 3000).

### Cytometry Time of Flight (CyTOF) analysis

One million single cells (tumor and tumor-associated immune cells) were resuspended in Maxpar PBS (Fluidigm, 201058). Dead cells were incubated with 0.75 µM Cisplatin for 5 min (Fluidigm, 201064), washed with Maxpar cell staining buffer (Fluidigm, 201068). FC receptors were blocked with TruStain FcX in 100 µL MaxPar cell staining buffer for 30 min at room temperature. Cells were then washed and stained with a cocktail of metal-conjugated antibodies for 30 min at room temperature and washed in a MaxPar cell staining buffer. Optimal concentrations were determined for each antibody by titration and the primary antibodies used are listed in **Supplementary Table 1**. Cells were resuspended and fixed in 1.6% paraformaldehyde prepared in MaxPar cell staining buffer for 20 minutes and washed in MaxPar cell staining buffer. Nuclei were labeled by incubating fixed cells in 1:4000 DNA intercalator (Fluidigm, 201192B) dissolved in MaxPar Fix and Perm Buffer (Fluidigm, 201067) for an hour at room temperature or overnight at 4°C. Following nuclear labeling, cells were washed once in MaxPar cell staining buffer and twice in MaxPar Water (Fluidigm, 201069). Samples were brought to 500,000 particulates per mL in MilliQ water containing 0.1x EQ beads (Fluidigm, 201078) and run in 450 μL injections on a CyTOF2 instrument.

CyTOF data was either analyzed using Cytobank online software or FlowJo. Cells were identified as events with high DNA (EQbeads served as negative control) and viable cells as events with high DNA and low Cisplatin. Once the viable cells were gated, CD45^+^ cells were identified as immune cells. These immune cells were downsampled to the number of immune cells present in MTD tumors for viSNE analysis, as MTD hosted the least number of immune cells. CD45^+^CD11b^+^ cells were identified as myeloid cells and CD45^+^CD3e^+^ cells as Lymphoid T cells. Myeloid cells further parsed into Neutrophils, Monocytes, and Dendritic (DCs) cells. The Cell-IDs of these cell subsets are: Neutrophils: Ly6C^low^Ly6G^high^, Monocytes: Ly6C^high^Ly6G^high^, NK cells: Nk1.1^+^, DCs: CD11c^high^ I-A-I-E^high^. T cells further parsed into CD4^+^ helper T cells and CD8a^+^ cytotoxic T cells. Activated T cells were identified as CD3e^+^CD44^+^ and regulatory cells as CD3e^+^CD25^+^. This gating strategy is illustrated in Fig. S1I.

### Cellular Indexing of Transcriptomes and Epitopes (CITE-seq) analysis

CITE-seq integrates cell surface protein and its transcriptome measurements as single-cell readouts using oligonucleotide based antibodies (Stoeckius et al., 2017). Antibodies used to phenotype different immune cells and their specific functional status are listed in **Supplementary Table 2**. Three different biological tumors were used for both the treatment conditions. A million immune cells were isolated using CD45 micro magnetic beads (Miltenyi Biotec, 130-052-301) from each tumor. Multiplexed with CITE-seq antibodies (CITE or ADT antibodies) and Cell hashing antibodies (HTOs, to identify samples), 20,000 cells were encapsulated using 10x Genomics Chromium single cell 3’ library and gel bead kit V3 (PN-1000092). On this chip, single cell was captured into emulsions with beads (GEM) to prepare cDNA libraries. Along with the primers for cDNA amplification PCR, primers for ADTs (CITE-seq antibodies) and the HTOs (HTOs 1 to 6, hashtag antibodies) amplification were included to increase the yield of libraries. Sequencing was performed at the IU Center for Medical Genomics for sequencing on Illumina NovaSeq 6000, S2, platform. The number of reads were 754 million and 380 million for gene expression and cell surface antibody expression respectively. The raw output from the sequencer was demultiplexed into sample specific mRNA, ADT, and HTO FASTQ files and were processed using Cell Ranger v3.1.

Seurat v3.0 (Stuart et al., 2019) R package was used for downstream multimodal analysis. Each sample was demultiplexed based on HTO hashtagging. High quality singlets were subset based on mitochondrial gene content less than 20% and gene expression RNA features between 200 - 8000 (Maier et al., 2020; Stuart et al., 2019). The gene expression matrix was normalized and highly variable features were selected for downstream analysis. Cells were downsampled (2356 per treatment condition) to have equal number of cells in either Vehicle or MCT-treated group for downstream analyses. Clusters formed from such quality controlled 2356 cells per treatment condition were referred to as Seurat Clusters. RNA assay and gene expression matrix were used for Principal Component Analysis (PCA) and Unified Manifold Approximation and Projection (UMAP). Clustered cells were further analyzed for differential gene expression, gene set enrichment (ssGSEA), trajectory analysis, and single-cell regulatory network inference analyses (SCENIC).

### CITE-seq gating strategy

After identifying singlets, *Ptprc* high and CD45 high cells were identified as immune cells. After downsampling (542 single cells per treatment condition), they were further parsed into CD11b^+^ myeloid cells and CD3e^+^ T cells. Then Ly6C^high^ Monocytes were gated out. Further, CD24^high^ F4/80^high^ cells were gated to identify CD11b^+^ and CD11c^+^ TAMs. On the other hand, CD24^high^ F4/80^low^ cells were gated to identify CD11b^+^ DCs and CD103^+^ DCs (together TADs). They were referred to as TADs and TAMs. The lymphoid cells with CD3e high expression further parsed into CD4 high and CD8a high cells. This gating strategy is illustrated in Fig. S2A and has been adopted from previous publications (Broz et al., 2014; Laoui et al., 2016).

### Dynverse trajectory inference analysis

Dynverse package (Saelens et al., 2019) was used for single-cell trajectory inference. All the myeloid clusters which shared transcriptional similarities with TADCs and TAMs (Seurat Clusters 0, 1, 3, 4, 7, 8, 10, 12, and 14) were selected as Dyno subclusters for trajectory inference. Cluster markers identified were: *S100a9*, *mt-Co3*, *Arg1*, *H2-Ab1*, *Cd74*, *Mrc1*, *Ccl8*, *Ly6c2*. These markers were selected as the start and end points of trajectories. Then, PAGA tree analysis function was used to infer the trajectory of relevant Seurat Clusters.

### SCENIC analysis

Single-Cell Regulatory Network Inference and Clustering (SCENIC) version 1.1.2.1 (Aibar et al., 2017; Van de Sande et al., 2020) was used for building the regulons and targetomes/gene regulatory networks (GRN). The analysis was performed using R software. For efficient analysis, the entire data set was down sampled to 1000 cells. For building the GRN, transcriptional factors and their top 10 potential gene targets were predicted via regression-based network inference. Putative regulatory regions of these positively regulated targets were searched and the enriched motifs were identified. The motifs with a NES > 3 were considered significant and positive regulators were chosen with pearson 0.03 (these were the values set by default). In some instances, for separating regulons with greater confidence in predicting motifs, NES ≥ 4 or 5 was chosen. We analyzed the data in three different iterations (1) All the myeloid cells pooled together: for **Fig. 7C**, split by Seurat Clusters (used to generate heatmap in **Fig. 7C**) (3) Individual Seurat Clusters (**Fig. 7D**). Each dot in the scatter plot represents a specific NES. We realized that multiple motifs of a regulon could have a NES and hence not to clutter the plot, we chose to represent each regulon with its highest NES (represented along the Y axis) with a dot. The scatter plots were generated after modifying the entire gene sets listed in their respective Supplementary Data Tables. For Cluster 0 *Stat1* targetome analysis: (1) The entire gene list is presented in **Supplementary Table 10**. (2) The scatter plot in **Fig. 8A**, represents Stat1 gene targets with NES > 3 and DEG in MCT with p < 0.05.

### T cell proliferation assay

After splenocytes were isolated as mentioned above (Single-cell isolation), Pan T Cell Isolation Kit II, mouse (Miltenyi Biotec, 130-095-130 or BioLegend, 480024) was used to isolate T cells. Using Fluorophore Assisted Cell Sorting (FACS), CD11c^+^ cells were isolated from CD45^+^ cells. CD45^+^ cells were isolated using magnetic beads (Miltenyi Biotec, 130-052-301). Using anti-PE Nanobeads (BioLegend, 480080) against PE fluorophore conjugated flow antibodies, CXCL16^+^ (BD Biosciences, 566740) cells were isolated from MCT-treated tumors. T cells were stained with CFSE stain following manufacturer’s instructions. T cells and CD11c^+^ cells were cultured in 1:1 ratio with IL2 in the heat-inactivated RPMI medium for 72 hrs. As experimental controls, T cells were either cultured alone or with T cell activation beads (Dynabeads™ Mouse T-Activator CD3/CD28 for T-Cell Expansion and Activation, Thermo Fisher Scientific, 11456D). After 72 hrs of culture, the cells were stained with CD3e, CD8a, and CD4 anti-mouse antibodies before proceeding to Flow Cytometry.

### T cell suppression assay

After splenocytes were isolated as mentioned above (Single-cell isolation), Pan T Cell Isolation Kit II, mouse (Miltenyi Biotec, 130-095-130 or BioLegend, 480024) was used to isolate T cells. Using Fluorophore Assisted Cell Sorting (FACS), CD11b^+^ Ccr1^high^ and CD11b^+^ Ccr1^low^ cells were sorted from CD45^+^ cells. CD45^+^ cells were isolated using magnetic beads (Miltenyi Biotec, 130-052-301) and APC or APC/Cy7 anti-mouse CD11b (BioLegend, 101212 or 101225) PE anti-mouse Ccr1 (BioLegend, 152507) markers were used for FACS sorting. T cells were stained with CFSE stain following manufacturer’s instructions. T cells with T cell activation beads and either CD11b^+^ Ccr1^high^ or CD11b^+^ Ccr1^low^ cells were cultured in 1:1 ratio in the heat-inactivated RPMI medium supplemented with IL2 for 72 hrs. As experimental controls, T cells were either cultured alone or with T cell activation beads (Dynabeads™ Mouse T-Activator CD3/CD28 for T-Cell Expansion and Activation, Thermo Fisher Scientific, 11456D). After 72 hrs of culture, the cells were stained with CD3e (BioLegend, 100319), CD8a (BioLegend, 100762), CD4 (Biolegend, 100410), PD-1 (BioLegend, 135217), and CXCR6 (BioLegend, 151120) anti-mouse antibodies and proceeded for Spectral Flow Cytometry (Cytek NL 3000).

### Adoptive Transfer of CD45.1^+^ cells

Tibiae, Fibulae, and Femurs were isolated from CD45.1^+^ B6 mice into heat-inactivated RPMI medium. The bones are then crushed with a pestle and washed and passed through a 40 µm cell strainer (VWR, 76327-098). RBC lysed and dead cells were separated (using dead cell removal kit, Miltenyi Biotec, 130-090-101). The live cells were then resuspended in a sort buffer to inject 1 million cells per B6 MMTV-PyMT mouse retro-orbitally.

### Intratumoral Injection of IgG or CXCL16 neutralization antibody

10ng of IgG or CXCL16 neutralizing antibody was dissolved in diluent to make upto 10 µL of the antibody cocktail and delivered at a constant rate (5µL per min) into the tumor using a Harvard Syringe Pump at a 45° angle. The tumors more than 10*10 mm in size were given 20 µL of the antibody. The tumors less than 5*6 or greater than 12*13 mm^3^ in size were not included in the study. For a combinatorial PD-1 inhibitor with CXCL16NAb (Fig.4 and Fig.6), 30ng in 10µL of CXCL16 was given per dose per tumor.

### shRNA lentiviral transfection assay

Bacterial stocks for pLKO or shRNA mouse *Stat1*(Bacterial Glycerol Stock Sequence (same as Construct 1) 1 # CCGGGCTGCCTATGATGTCTCGTTTCTCGAGAAACGAGACATCATAGGCAGCTTTTTG, Bacterial Glycerol Stock Sequence (same as Construct 2) 2# CCGGGCTGTTACTTTCCCAGATATTCTCGAGAATATCTGGGAAAGTAACAGCTTTTTG) constructs were purchased from Sigma Aldrich. The plasmids were packaged into Lentiviral particles using the 2rd generation packaging plasmid (Addgene). 10 ml of virus containing media were concentrated with Lenti-X concentrator (Takara Bio, 631231) to a titer sufficient to decrease pStat1 levels by 25% (Mean pStat1+ CD11b+ cells: pLKO: 100, Construct 1# 75, Construct 2# 75). MCT-treated CD11b^+^ cells were isolated from tumors using MojoSort Mouse CD11b Selection Kit (BioLegend, 480109). For transfecting every million primary CD11b^+^ cells, 1ml of the viral concentrate was used along with 5 μg/mL of polybrene. The cells were transfected for 72 hrs prior to analysis by flow cytometry. For flow cytometry analysis, the cells (post transfection) were washed with a cell staining buffer, blocked for 20 min on ice in dark and stained with APC/Cy7 CD11b antibody (BioLegend, 101225) and PE anti-mouse PD-L1 (BioLegend, 124308) flow antibodies and incubated for 20 min in dark on ice. The cells were further fixed and permeated using a phospho permeable buffer. Later, cells were washed, stained with Alexa Fluor 488 pStat1 (Ser 727) (Biolegend, 686410) for 20 min, washed, and resuspended in a cell staining buffer before analysing on the flow cytometer.

### Multiple gene association data analysis

For multiple gene association analysis, GEPIA web server was used. GEPIA uses RNA-seq data from TCGA and GTex projects. Here we compared *Stat1* and *Cd274* correlation between healthy breast, blood, ovarian, skin, and lung tissues and cancerous tissues (BRCA, CHOL, LAML, HNSC, OV, SKCM, SARC). We observed that *Stat1* and *Cd274* correlate stronger in cancerous conditions (R = 0.71) than in healthy states (R = 0.18) (Fig. S5G).

### Tissue processing

For formalin-fixed, paraffin-embedded tissues (FFPE), the tumor tissues covered in cassettes were incubated in 10% nebulized buffered formalin overnight at room temperature. The tissue cassettes were then incubated in 70% ethanol for a week before being processed in the Leica Tissue Processor using an 8 hour program specific for fat rich tissues. To perform staining, 4µm thick tissue ribbons were sectioned using Leica Tissue Microtome. These tissue ribbons were then adhered to SuperFrost slides (VWR, 48311-703) and dried overnight before being processed for staining.

### Tissue Immunohistochemistry (IHC)

FFPE tissue sections were deparaffinized by incubation in a 60°C oven for 30 min and in Xylene for 10 min. Hydration was achieved by sequentially immersing slides for 10 min each, in staining dishes containing 100%, 95%, 80% ethanol, and ddH_2_O. Antigens were unmasked by boiling slides in 1x Sodium citrate solution (10mM Sodium citrate, pH 6.0) in a pressure cooker for approximately 20 min. After antigen retrieval, tissues were blocked in 3% H_2_O_2_ in phosphate buffered saline (PBS) for one hour at room temperature to quench endogenous peroxidase. To quench nonspecific antibody binding, the tissue sections were blocked in horse serum for an hour. Tissues were then stained with primary antibodies overnight in a humidity chamber at 4℃. Following overnight incubation, the tissues were washed in TBST three times 30 min each before an hour of incubation with secondary antibodies. The Vectastain Elite ABC-HRP Kit RTU (Vector Laboratories, PK-7200) was used to develop the peroxidase before incubating with the ImmPACT-DAB chromogen. The primary antibodies used were Ki-67 (clone D3B5, Cell Signaling Technology, CST 12202S) and anti-CD3 (Abcam, ab 16669).

### Tissue Immunofluorescence (IF)

FFPE tissue sections were first incubated in a 60°C oven for 30 min and washed twice with Xylene for 5 min. Hydration was achieved by sequentially immersing slides, for 10 min each in 100%, 95%, and 80% ethanol. Then the slides were washed in ddH_2_O before proceeding to the antigen retrieval step. Antigens were unmasked by boiling slides in 1x Sodium citrate solution (10mM Sodium citrate, pH 6.0) in a pressure cooker for approximately 20 min (first 10 min without weight at 350℃ and next 10 min with weight at 240℃). After cooling to room temperature, slides were washed 3 times with PBS in vertical staining jars. Dewaxed specimens were blocked by incubation with NaOH and 4.5% H_2_O_2_ for an hour. Then the slides were incubated with Odyssey blocking buffer (LI-COR Biosciences, 927-40150) or 2.5% Donkey Serum in PBS for 60 mins by applying the buffer to slides as a droplet at room temperature; evaporation was minimized by using a humidity chamber. Slides were incubated overnight with primary antibodies (both fluorophore conjugated and unconjugated). The unconjugated primaries were further incubated with secondaries for two hours before washing three times with PBS, 5 min each. Finally, slides were incubated with Hoechst 33342 (1 mg/ml) in Odyssey blocking buffer for 30 min and washed three times in PBS, 5 min each. A set of immune cell markers and proteins of interest were used to stain breast tissues (and Spleen for positive control when appropriate) which are treated either with PBS, MCT, MTD, anti-PD-1, CXCL16 NAb, Stat1 inhibitor or a combination of either of these. They are: anti-Iba1 (dilution 1:2000, Abcam ab5076), Phosphorylated STAT1 Stat1 - Ser 727(dilution 1:200, BioLegend 686405), anti-CD3 (dilution 1:500, Abcam,ab 16669), anti-CD8 (dilution, CST 98941S), anti-PD-L1 (dilution 1:500, CST 64988S), and anti-CXCL16 (dilution 1:500, R&D Systems MAB503-100).

### Statistical analysis

All common quantitative data were analyzed and plotted as box plots or survival curves using the R program. Seurat, ssGSEA, Dynverse, and SCENIC analyses were also performed in the R program as described in each dedicated section. Flow cytometry and CyTOF data was analysed using FlowJo software. viSNE analyses of CyTOF were performed using Cytobank. For quantitative image analysis, positively stained cells (myeloid, lymphoid, and their colocalization with markers such as PD-L1 and CXCL16) or Ki67 high to low cells were counted manually in a given field of view using ImageJ.

## DATA AND CODE AVAILABILITY

All CITE-sequencing Data has been deposited at GEO (accession code GSE158888, reviewer access token: qzovmuyqdvufhmb). All raw data will be made available publicly upon acceptance for publication. The R code for data analysis will available upon publication at GitHub.

---

**Figure S1 related to Figure 1.**
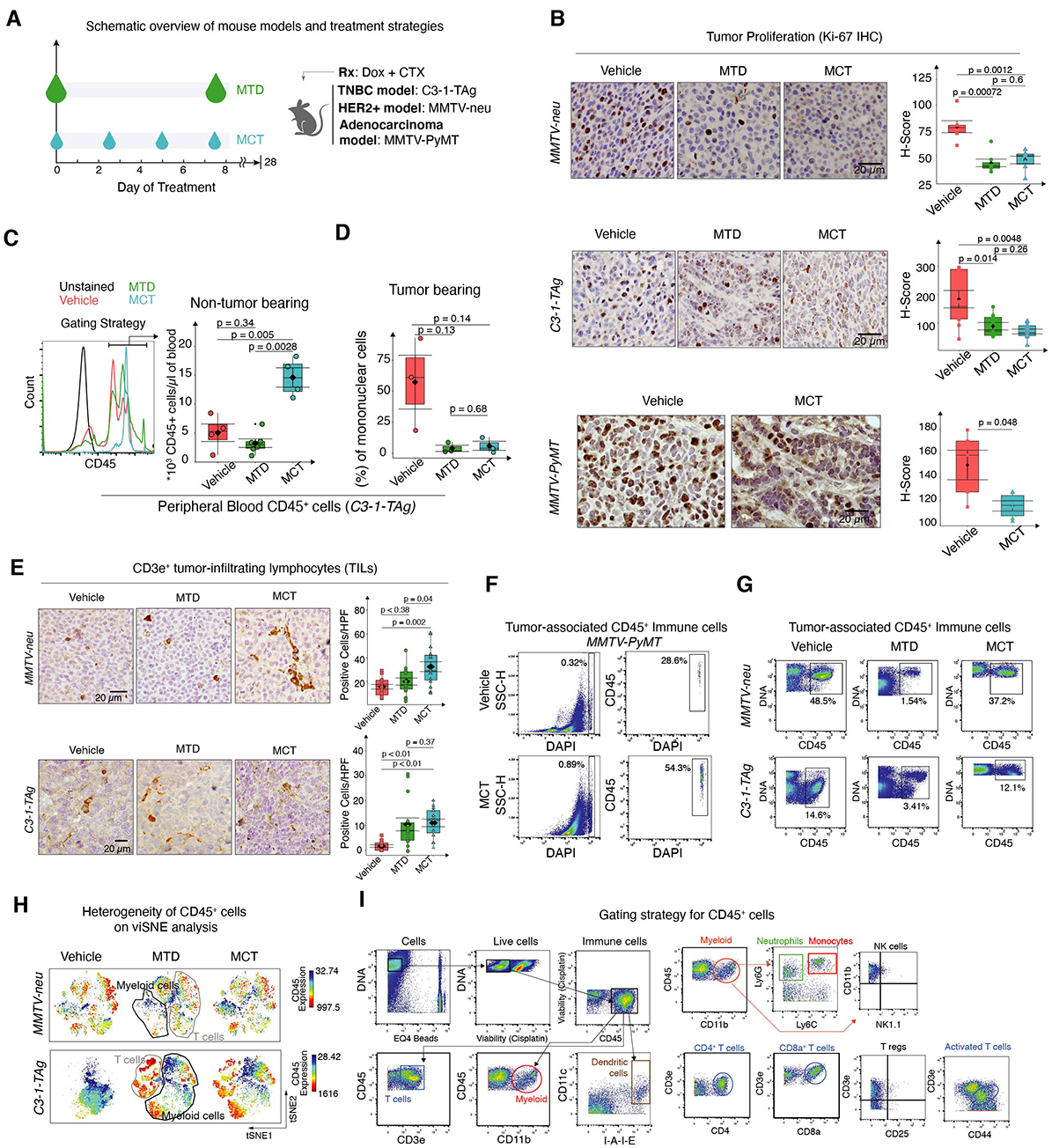
(A) Schematic overview shows the MTD and MCT treatment strategies. (B) Representative IHC images of MMTV-neu (top) and C3-1-TAg (middle) and MMTV-PyMT (bottom) breast tumor tissues stained with Ki-67 and box plots quantify H-Score (n = 6 to 9 slides per group, Scale bar = 20 *µ*m). (C) Flow cytometry gating strategy to identify CD45^+^ cells (left) and box plots (right) show quantification of CD45^+^ cells in peripheral blood of non-tumor bearing mice. (D) Box plot quantify CD45^+^ cells in peripheral blood of tumor bearing C3-1-TAg mice. (E) Representative IHC images of MMTV-neu (top) and C3-1-TAg (bottom) stained with CD3 and box plots quantify CD3^+^ cells per high power field (HPF) (n = 15 slides per group). (F) Spectral flow gating strategy to identify tumor-associated CD45^+^ immune cells in MMTV-PyMT mice. (G) CyTOF gating strategy to identify tumor-associated CD45^+^ immune cells in MMTV-neu and C3-1-TAg mice. (H) t-SNE plots show the tumor-associated immune cell distribution an their CD45 expression. (I) CyTOF data biaxial plots show the gating strategy to identify different tumor-associated live immune cell populations. Data in box plots represent Mean ± SEM, Diamonds Means, Black lines Medians, and T-test p-values. n = 2 for CyTOF analysis and n ≥ 3 (biological replicates) for the rest.

**Figure S2 related to Figure 2.**
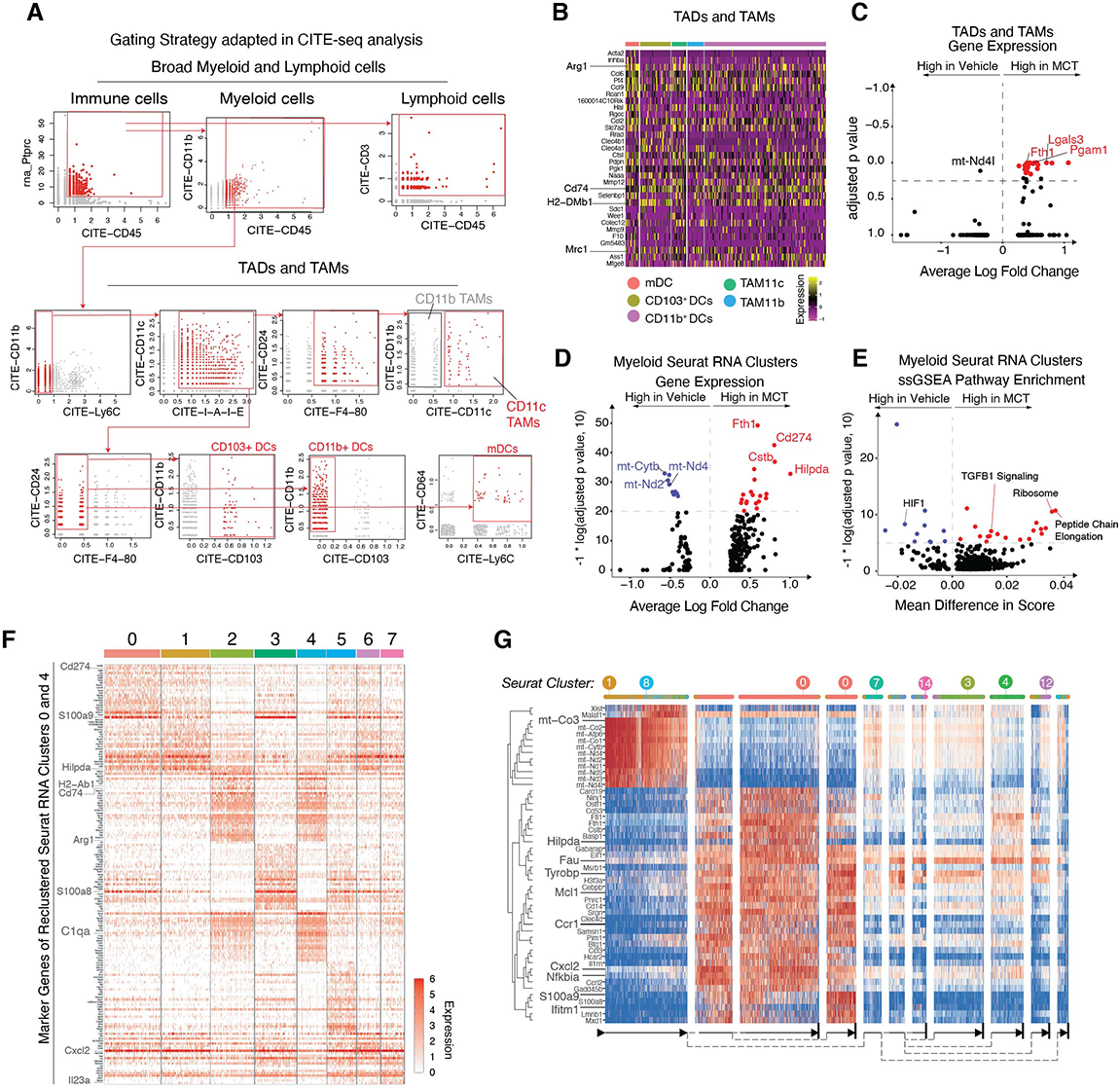
CITE-seq reveals distinct myeloid subsets coexist in MCT TME. (A) Gating strategy adopted in CITE-seq analysis. (B) Heatmap shows OGE between different TADs and TAMs. (C) Volcano plot of DEGs between MCT and Vehicle treated TADs and TA Ms. Volcano plots of (D) DEGs and (E) DEGPs between MCT and Vehicle treated Seurat Myeloid clusters from Fig. 2C. (F) Heatmap shows top DEGs between clusters identified in Fig. 2E. (G) Heatmap shows expression of marker genes of predicted trajectory paths.

**Figure S3 related to Figure 3.**
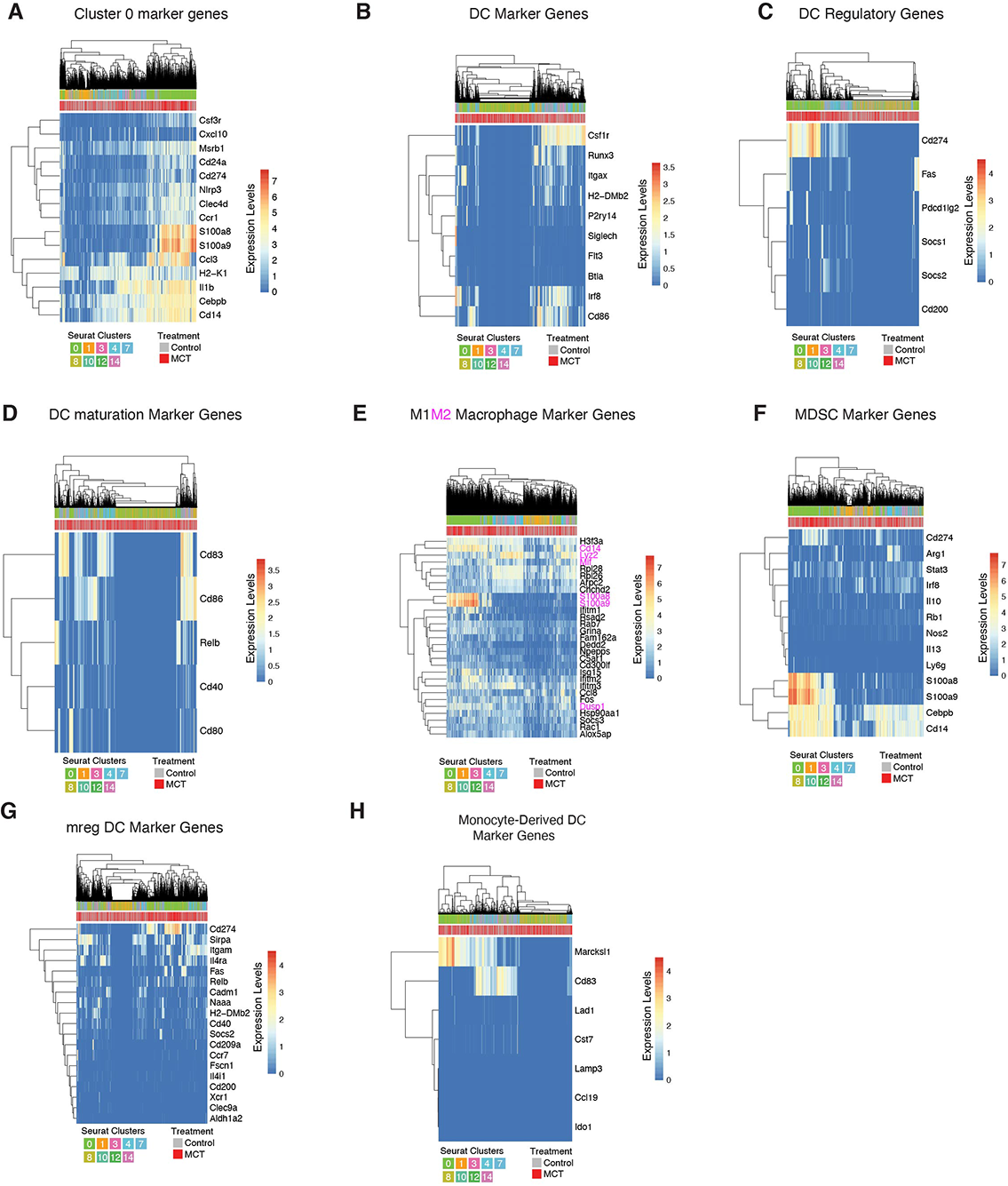
Transcriptional Profile of Cluster O. Heatmaps show (A) Cluster O DEGs (B) DC (C) DC regulatory (D) DC maturation (E) M1M2 macrophage (F) MDSC (G) mature regulatory (mreg) DC (H) Monocyte-Derived DC marker gene expression by Seurat Clusters 0, 1, 3, 4, 7, 8, 10, 12, and 14.

**Figure S4 related to Figure 4.**
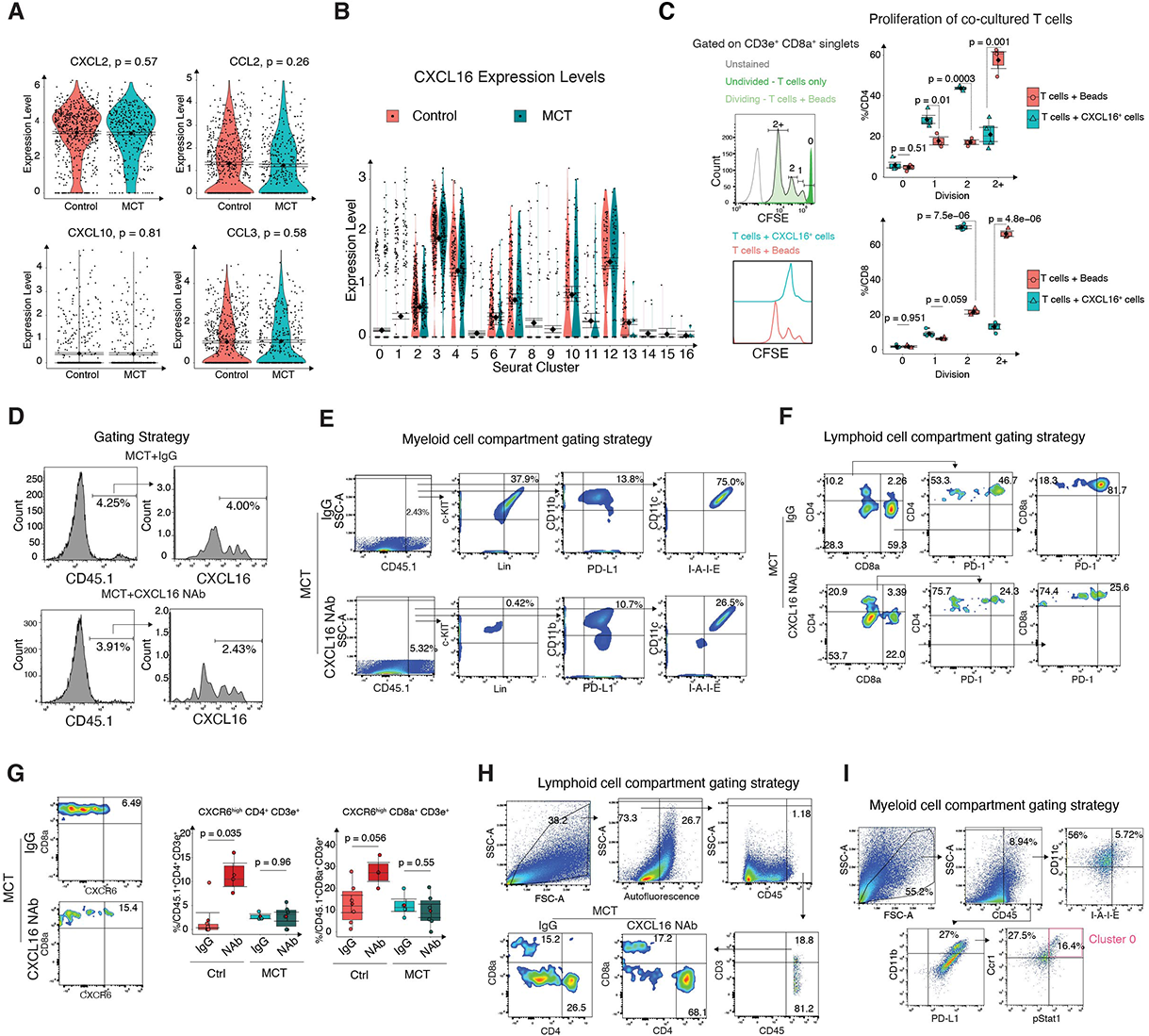
MCT-induced CXCL16 mediates intratumoral immune dynamics and ICB efficacy. (A) Violin plots compare the expression of different cytokine genes by DC clusters between treatment groups (B) Violin plot compares expression of CXCL16 between Seurat clusters. (C) CFSE dilution T cell proliferation assay: Gating strategy (left) indicates various division phases on positive control (T cell activation beads) CD8+ T cells (top), histogram outlines compare the division phases (CFSE levels) between T cells from CXCL16 and bead co-culture (bottom). (D) Gating strategy to identify CXCL16+ CD45.1 + cells comparing between lgG and NAb groups of MCT-treated mice. Gating strategy to identify (E) c-kit+ Lin+, CD11b+PD-L1+, CD11c+ I-A-1-E+ cells (F) CD4+, CD8a+, CD4+ PD-1+, CD8a+ PD-1+ (G) CXCR6+ CD4 and CD8a+ T cells (left) and (G) quantification of CXCR6+ T cells (right). Gating strategy to identify (H) CD45+ CD8+ and CD4+ T cells and (I) CD11c+ I-A-1-E+ cells and Cluster O cells in lgG and NAb traeated MCT primed C3-1-TAg mice. Data in box plots represents Mean ± SEM, Diamonds Means, Black lines medians, and T-test p-values.

**Supplementary Figure 5 realted to Figure 5.**
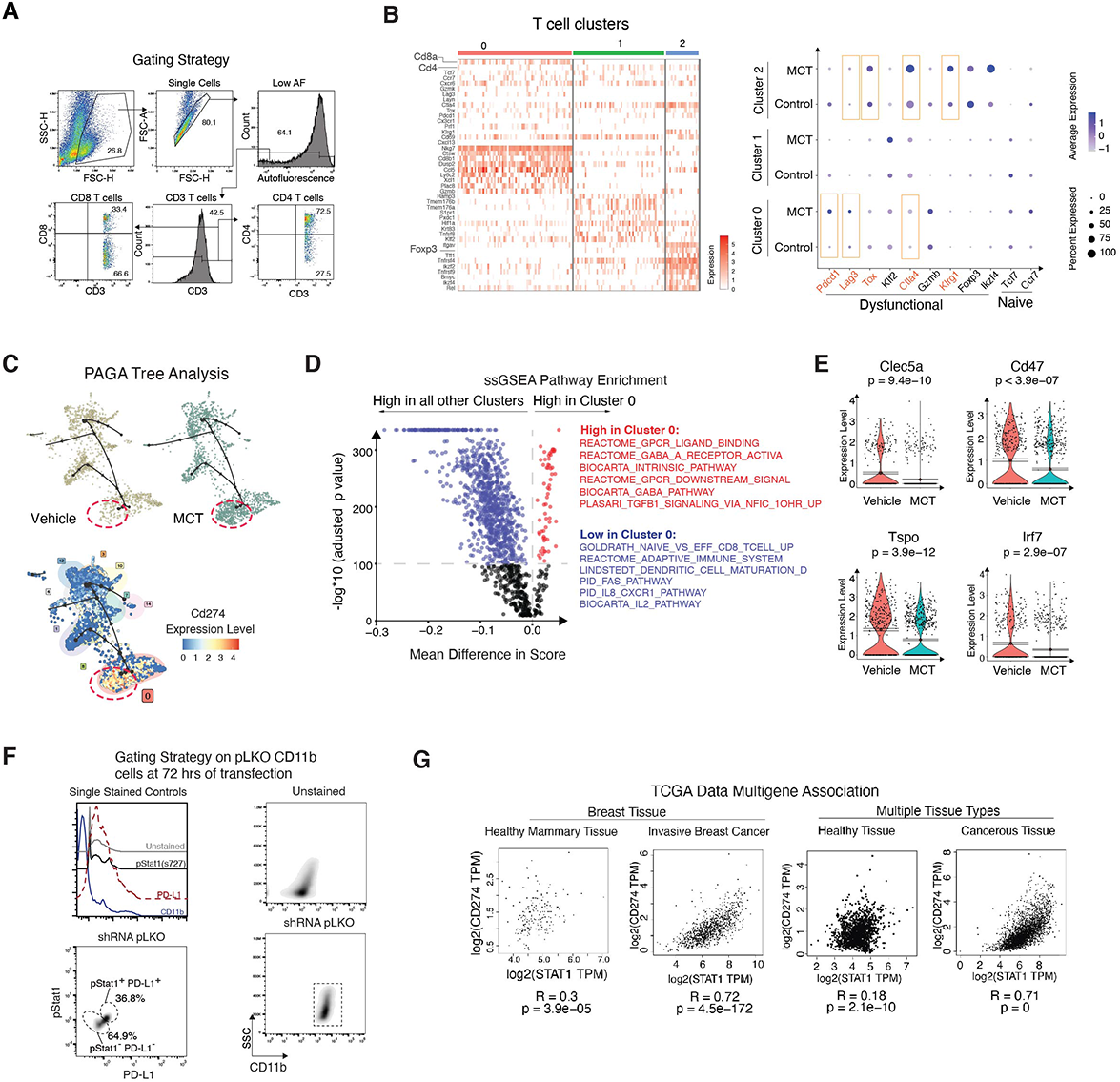
STAT1 regulates PD-L1 expression in Cluster O cells and influences T cell activation status. (A) Gating strategy to identify CD8 and CD4 T cells in co-culture shown in Fig. 5A. (B) Heatmap (left) shows identity of different Cd3e high T cells. Dotplot (right) split by treatment group to show percentage of cells expressing various functional and differentially expressed genes. (C) PAGA trajectory trees (left, top) show Seurat Clusters (encircled are Cluster 0) split by treatment group and expression of Cd274 (left, bottom). (D) Volcano plot shows enriched DEGP from ssGSEA in Cluster O cells over other Seurat Clusters. (E) Violin plots show expression of Clec5a, Cd47, Tspo, and lrf7 genes by Cluster O cells (each dot is a cell and cells are pooled from n = 3 biological replicates). (F) Flow cytometry histograms show the individual antibody levels on single stained pLKO transfected cells (left, top). Biaxial plots show the gating strategy adopted to identify proportions of pStat1^+^ and PD-L1^+^ cells after shRNA transfection (left, bottom). Biaxial density plots comapre CD11b levels in unstained and stained shRNA pLKO *in vitro* cultures. (G) Scatter plots show association between Stat1 and Cd274 across healthy mammary and invasive breast cancer tissue and multiple healthy tissues and their respective cancerous tissues. The plots were generated using Gepia web server and R is Spearman’s correlation.

**Figure S6 related to Figure 6.**
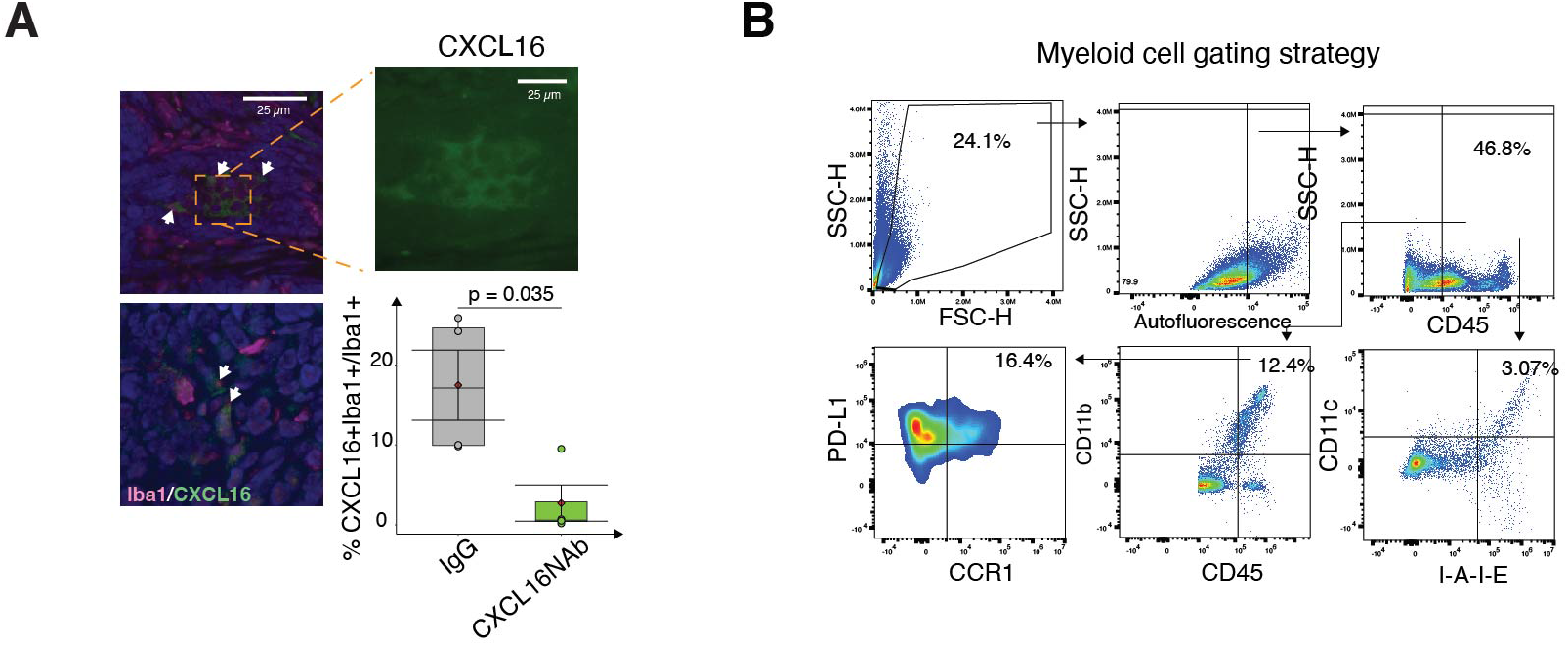
(A) Representative IF images show and box plots quantify the CXCL16+ myeloid cells upon treating with CXCL16 NAb (Regimen 2). (B) Flow plots show gating strategy to identify myeloid cells. Data in box plots represents Mean ± SEM, Diamonds Means, Black lines medians, and T-test p-values.

